# Information theoretic evidence for layer- and frequency-specific changes in cortical information processing under anesthesia

**DOI:** 10.1101/2022.07.15.500162

**Authors:** Edoardo Pinzuti, Patricia Wollstadt, Oliver Tüscher, Michael Wibral

## Abstract

Nature relies on highly distributed computation for the processing of information in nervous systems across the entire animal kingdom. Such distributed computation can be more easily understood if decomposed into the three elementary components of information processing, i.e. storage, transfer and modification, and rigorous information theoretic measures for these components exist. However, the distributed computation is often also linked to neural dynamics exhibiting distinct rhythms. Thus, it would be beneficial to associate the above components of information processing with distinct rhythmic processes where possible. Here we focus on the storage of information in neural dynamics and introduce a novel spectrally-resolved measure of active information storage (AIS). Drawing on intracortical recordings of neural activity under anesthesia before and after loss of consciousness (LOC) we show that anesthesia-related modulation of AIS is highly specific to different frequency bands and that these frequency-specific effects differ across cortical layers and brain regions.

We found that in the high/low gamma band the effects of anesthesia result in AIS modulation only in the supergranular layers, while in the alpha/beta band the strongest decrease in AIS can be seen at infragranular layers. Finally, we show that the increase of spectral power at multiple frequencies, in particular at alpha and delta bands in frontal areas, that is often observed during LOC (‘anteriorization’) also impacts local information processing – but in a frequency specific way: Increases in isoflurane concentration induced a decrease in AIS in the alpha frequencies, while they increased AIS in the delta frequency range *<* 2Hz. Thus, the analysis of spectrally-resolved AIS provides valuable additional insights into changes in cortical information processing under anaesthesia.

**Author Summary:** While describing information processing in digital computers is somewhat straightforward and accessible (e.g. how much information is stored in a hard disk or which modification of information a CPU is executing), quantifying the widely distributed information processing in a biological neural system is much more challenging. In neural systems separating the components of distributed information processing - information transfer, storage and modification - helps with this task, but requires accurate mathematical definitions of these components of information processing. These definitions of distributed information processing quantities have become available only very recently. Of the three component processes mentioned above information storage, in particular, has been used with great success to analyze information processing in swarms, and to evolve, and optimize artificial information processing systems. The analysis of information storage has also already proven to be useful for the analysis of biological neural systems. Since in such systems, information processing seems to be often carried out by rhythmic neural activity with different frequencies, a measure of the frequency-specific components of the active information storage is needed. Here introduce such a measure and study how isoflurane anesthesia affects the local information processing in the ferret prefrontal and primary visual areas around loss of consciousness. We found that the modulation of active information storage by isoflurane is specific to frequency, layers and area, and that the analysis of frequency-specific active information storage provides insights not captured by more traditional descriptions of neural activity.

## Introduction

Biological systems must process information about their environment and their internal states in order to survive. Many biological systems have evolved specialized areas where such information processing is particularly evident. Prime examples are the central nervous systems of many animals and the human brain in particular. Taking inspiration from such systems, humans have developed biologically-inspired, artificial information processing systems, such as artificial neural networks, to solve a variety of tasks. Artificial neural networks and their biological sources of inspiration share an important property—they perform highly distributed information processing in which fundamental information-processing operations such as storing, transferring and modifying information are both, highly distributed and co-located at almost all computational elements. The computational elements making up biological and artificial neural networks, for example, are neurons, where each neuron’s activity can simultaneously serve the storage, transfer and modification of information. This lack of specialization and high degree of distribution separates such information processing systems from classical digital architectures (like a household PC) where the fundamental information processing operations are much more spatially separated and carried out by dedicated subsystems. While the highly distributed information processing certainly adds to the performance of artificial and biological neural networks on certain tasks, it also poses a formidable challenge to understand how such a system functions.

A powerful approach to describe and understand computation in systems such as biological or artificial neural networks is information theory, which introduces measures of information transfer, storage and modification [1–5]. The proposed measures are well-suited to investigate the function of artificial information processing systems, and have successfully been applied to biological neural systems [6–9]. However, in its original form, the framework neglects a central aspect of information processing in biological neural networks, namely the frequently displayed highly rhythmic activity when performing a computation. To understand those systems better and to build a bridge between information processing and their biophysical dynamics, it would therefore be beneficial to link the components of information processing to specific neural rhythms.

We have recently presented such a link for the case of information transfer in [10], and have provided results that challenged some long-held *ad hoc* beliefs about the relationship of brain rhythms and information transfer. In the present work, we extend this approach to information storage. In particular we focus on the *active storage*, where the information storage is actively in use for a computation in the dynamic of the neural activity (for differences with passive storage, e.g synaptic gain changes, see [11]). A measure of this kind of storage is the *active information storage* (AIS) [3, 7], which quantifies the amount of information in the present samples of a process (“currently active”) that is predictable from its past value. The AIS measure is closely linked to the transfer entropy (TE) [1]: the TE quantifies information transferred from a source process to the current value of a target process, in the context of the target process’ own past. Hence, AIS and TE together reveal the sources of information which contribute to prediction of the target process’ next outcome (either, information actively stored in the processes’ own past, or additional information being transferred from another process) [7].

The importance of understanding how neurons and neural systems store information when studying neural information processing has been outlined already in [11] and later by the work of [3, 7, 12]. AIS as a measure of information storage has been successfully applied in magnetoencephalographic (MEG) recordings to test, for example, predictive coding theory [13] or to provide better understanding of the information processing in people affected with the Autism spectrum disorder (ASD) [6, 14]. In local field potential (LFP), [8] found an increased AIS measure as a function of anesthesia (isoflurane) concentrations in two ferrets recordings, at prefrontal (PFC) and visual cortical (V1) sites. Anesthetic agents such as isoflurane are known to affect the frequency spectrum throughout the cortex [15] and at laminar level [15, 16]. In [15] it was shown that the effect of isoflurane on neural oscillatory activity is not only frequency-specific but also related to the computational property of the area, being different between different areas of the cortex (PFC or V1) or between different layers (deep laminar or infragranular layers, granular layers, and superficial or supragranular ones).Similarly, [16] reported highly specific effects of isoflurane on laminar frequency data.

Even though the effect of anesthesia on brain rhythms is known, due to a lack of a suitable method, all attempts to link the AIS with the rhythmic activity in different frequency bands were only indirect and through correlation analysis [6, 8, 14]. Hence, it seems beneficial to have also a spectrally-resolved AIS to directly investigate effects, for example, of isoflurane agents on brain rhythms and thus on neural information processing. We here present such a method, which is able to quantify AIS in a spectrally-resolved fashion. We apply this method to laminar recordings from two areas of the ferret cortex (PFC and V1) under different levels of anesthesia, to investigate how different frequency bands contribute to information storage under anesthesia. We hypothesised that due to the different computational properties of the layers [17] (either deep or superficial), the frequency-resolved AIS will show a heterogeneity of relevant frequency bands across frequencies and recording sites. On the other hand, low frequency oscillations (1HZ) caused by higher levels of anesthesia, which are a mark for loss of consciousness (LOC) [18], could have a coherent effect across cortical layers. We apply our AIS measure to disentangle these different effects and evaluate how isoflurane affects the AIS measure across anesthesia levels and cortical areas by means of Bayesian Regression.

## Materials and Method

In this section, we first clarify the purpose and application of the proposed method. Second, we introduce the information theoretic preliminaries together with the AIS measure, and the corresponding notation. Central to our method is the creation of frequency-specific surrogate data, for which we summarize the technical background.

Here, we outline only the crucial properties of the Maximal Overlap Discrete Wavelet Transform (MODWT), while a more detailed description can be found in [19, 20]. Finally, we present the core algorithm to identify frequency-specific AIS.

## Background

### Problem Statement and Analysis Setting

The aim of the proposed method is to determine whether there is statistically significant active information storage generated by one or more frequencies. Our method can be implemented after a significant AIS has been determined in the time domain, e.g. as computed by the AIS algorithm in [21], in order to provide a perspective on this novel spectrally-resolved AIS.

### Technical Background: Active information storage (AIS)

We assume that a stochastic process 𝒴 recorded from a system (e.g cortical or layers sites), can be treated as a realizations *y*_*t*_ of random variables *Y*_*t*_ that form a random process **Y** = {*Y*_1_…, *Y*_*t*_, …, *Y*_*N*_}, describing the system dynamics. Then, *AIS* is defined as the (differential) mutual information between the future of a signal and its immediate past state [3, 7, 22]:

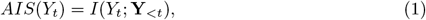

where *Y* is a random process with present value *Y*_*t*_, and past state 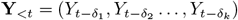, with *δ*_*i*_ = *i*Δ_*t*_, where Δ_*t*_ is the sampling interval of the process observation, and *δ*_1_ ≤ *δ*_*i*_ ≤ *δ*_*k*_. **Y**_*<t*_ is a vector of random variables chosen from **Y** from the past of the current time point *t*. The collection, or vector, **Y**_*<t*_ captures the underlying dynamic of the system 𝒴 and can be seen as a state space reconstruction, for details see [3, 23]. We here employed a recently proposed non-uniform embedding algorithm from the IDTxl toolboox [21] to properly construct the nonuniform embedding of *Y* time-series [24, 25]. This algorithm also yields approximations for parameters like *δ* and *k*. Thus, the AIS estimates how much information can be *predicted* by the next measurements of the process by examining its paste state [3]. In processes that either produce little information (low entropy) or that are highly unpredictable, the AIS is low, whereas processes that are predictable but visit many different states with equal probabilities [7], exhibit high AIS [7, 9].

### Technical Background: Maximum Overlap Discrete Wavelet Transform

Our method is based on the creation of suitable surrogate data for use in a statistical test. Many methods exist for surrogate data creation, each with its own limitations and advantages (see [26] for a review). Among these, wavelet-based methods allow to create the needed frequency-specific surrogate data through randomization of the wavelet coefficients [27]. In particular, wavelet-based surrogates that preserve the local mean and the variance of the data were introduced by [28]. Similarly to [29], we employ the Maximal Overlap Discrete Wavelet Transform (MODWT), to transform the data in the wavelet domain. The MODWT is well defined for time-series of any sample size and produces wavelet coefficients and spectra unaffected by the transformation. [29].

The MODWT of a time-series *X* = (*X*_0_, …, *X*_*N−*1_) of *J*_0_ levels, where *J*_0_ is a positive integer, consists of *J*_0_ + 1 vectors: *J*_0_ vectors of wavelet coefficients 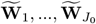 and an additional vector 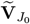 of scaling coefficients, all with dimension *N* (our exposition of the MODWT closely follows that of [19], pages 159-205). The coefficients of 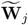, and 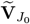, are obtained by filtering *X*, namely:

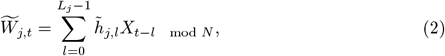

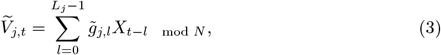

where 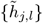 and 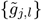 are the *j*th level MODWT wavelet and scaling filters, with *l* = 1, …, *L* being the length on the filter and *L*_*j*_ = (2^*j*^ − 1)(*L* − 1) + 1. We can write the above in matrix notation as:

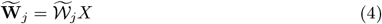

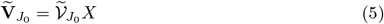

where each row of the *N* × *N* matrix of 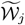 has values denoted by 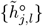, while 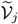 has values denoted by 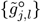, where 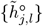 and 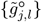 are the periodization of 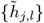 and 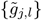 to circular filter of length *N* [19]. Thus, the MODWT treats *X* as if it were periodic, such periodic extension is known as “circular boundary condition” [19]. Finally, the time series *X* can be retrieved from its MODWT by [19]:

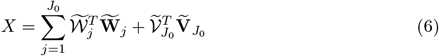

While, the coefficients 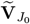 represent the unresolved scale [19, 29], and capture the long-term dynamics of *X*, the coefficients 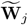 are associated with changes of the underlying dynamics, at a certain scale, over time. If *N* = 2^*J*^ and we set *J*_0_ = *J*, then a full decomposition is performed and the scale 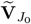 retains only the average constant of the data with all other information represented in the wavelet coefficients [29, 30]. Since in many applications a full decomposition is not necessary (e.g. the dynamic of a physical system is meaningful over a certain frequency range only), *J*_0_ can be set to any integer *J* ≤ ⌊ (log_2_(*N*)) ⌋so that the decomposition at any scale is shorter than the total length of the time series [31]. The selection of *J*_0_ determines the number of scales of resolution with the MODWT coefficients at a certain scale *j* related to the nominal frequency band |*f*| ∈ (1*/*2^*j*+1^, 1*/*2^*j*^) [19]. Moreover, given 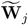 and 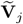, it is possible to reconstruct the time-series *X* through the inverse MODWT (IMODWT). If the coefficients are not modified, the IMODWT returns the original time-series *X* [19]. As shown in [10] the MODWT is a suitable and efficient method to create surrogate data as required by the current algorithm.

### Algorithm

To obtain a frequency-resolved AIS measure, our algorithm’s main idea is to create surrogate data, in which we destroy the AIS-relevant signal properties, i.e., the temporal order, in specific frequency bands. We then compare AIS estimates from the original data with estimates from the surrogate data, and establish via non-parametric statistical testing whether destroying specific frequency components led to a drop in AIS. This approach has has been successfully demonstrated in [10] to estimate frequency-specific TE and replaces approaches that use filtering or other preprocessing steps to estimate frequency-resolved measures, as these come with well known problems [10, 32]. As in [10], we here employed an invertible wavelet transform (maximum overlap discrete wavelet transform, MODWT) and a frequency- or scale-specific scrambling of the wavelet coefficients in time for surrogate data creation, keeping the original time-series always intact. With this method, we are also able to handle any potential bias introduced by the wavelet filtering of the surrogate data, yielding a more conservative analysis. Indeed, if the frequency-specific AIS measure should increase due to the scrambling of the wavelet coefficients, this will not result in a significant drop when statistically compared to the original AIS, and will thus not be mistaken for an effect.

### Implementation

Below, we will detail the algorithm for the measurement of frequency-specific AIS. As introduced above, we obtain this measure by creating surrogate data in which the temporal ordering of the signal has been destroyed for specific spectral components, by first transforming into the frequency domain, then scrambling wavelet coefficients and last transforming back to the time domain to obtain surrogate data. Overall this algorithm relies on five steps:

1. Perform a wavelet decomposition of the source time series through the MODWT to obtain a time-frequency representation of **Y** in *J*_0_ scales.
2. At the j^th^ scale of the MODWT decomposition shuffle the wavelet coefficients to destroy information carried by the scale (frequency band)
3. Apply the inverse wavelet transform, IMODWT, to get back the time representation of the time series
4. Compute the 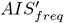 of the process **Y** .
  a. Repeat step 2 to 4 for a sufficiently high number of permutations to build a surrogate data distribution.
  b. Repeat step 1 to 4 for all *J*_0_ scales.
5. Test whether the original *AIS* is above the 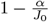 quantile of the surrogate-based distribution of 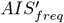 values at each scale, i.e. perform a significance test with respect to the surrogate-derived distribution.

The operations implemented in the five steps are illustrated in Fig. **??** and described in detail hereafter.

**Step 1:** The time-series is decomposed once into *J*_0_ scales through the MODWT (Fig. 1, A). As introduced in section *Maximum Overlap Discrete Wavelet Transform* this decomposition gives a set of coefficients 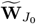 and an additional set of approximation coefficients 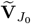. The latter is saved in this first step and utilized only in step 3, without any modification. Only the 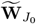coefficients at the *j*^*th*^ scale under analysis are subjected to step 2. The current implementation uses a Least Asymmetric Wavelet (LA) as mother wavelet of length 8 or 16, since both lengths showed to be robust against spectral leakage and do not relevantly suffer from boundary-coefficient limitations. [19, 28, 33].

**Fig 1.**
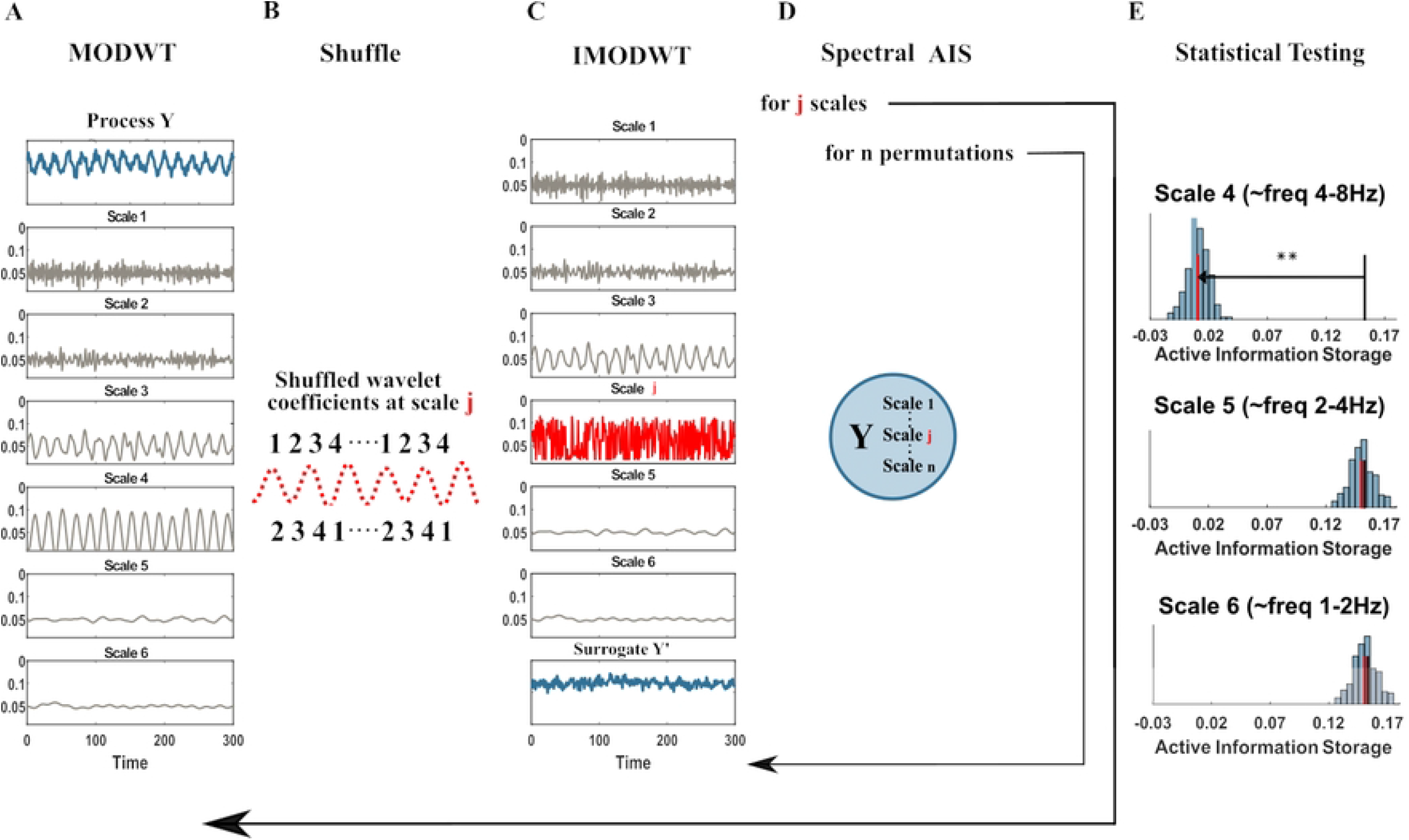
Spectral AIS algorithm pipeline. (A) The neural signal (blue) is converted to a time-frequency representation (grey) using the invertible maximum overlap discrete wavelet transform (MODWT). (B) At a frequency (wavelet scale) of interest in the source the wavelet coefficients are shuffled in time, destroying its internal dynamic. (C) The signal is recreated by the inverse MODWT. (D) The AIS for the original and many shuffled signals is computed. (E) A statistical tests determines whether the shuffling reduced the active information storage, indicating that the information storage was indeed encoded at the specific frequency. Each panel here shows the distribution of 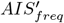values (vertical bars) obtained from surrogate data where the wavelet coefficients of the scale of interest were shuffled, the median of this distribution (red line), and the original AIS (black line). The analysis and the testing is repeated for all scales of interest (here 4,5,6) .

The creation of surrogate data for subsequent statistical testing comprises of the following steps 2 and 3.

**Step 2:** The frequency-specific active information storage of the process is destroyed by shuffling the 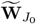wavelet coefficients one scale at a time. The *j* ^*th*^ scale under analysis is shuffled by randomly permuting the coefficients 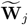 whereas all the other scales decomposed by the MODWT stay intact (Fig. 1, B, *j*^*th*^ scale in red). We implement two alternative methods for the creation of surrogate data: a Block permutation of the wavelet coefficients [27] and the Iterative Amplitude Adjustment Fourier Transform (IAAFT) [27, 29]. Since there is no canonical method of surrogate data creation and in many cases the employment of one method over another depends on the specific analysis carried out by the user.

**Step 3:** The unchanged set of coefficients, 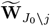, the unchanged 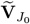’s, and the permuted coefficients at scale *j* 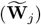 are submitted to the IMODWT, to reconstruct the surrogate process signal, **Y**′, in the time-domain (Fig. 1, C). This step is identical for both of the implemented surrogate-data creation methods: Block permutation of the wavelet coefficients and IAAFT. The reconstructed process **Y**′ (*process surrogate*) differs from the process **Y** only on the shuffled *j*^*th*^ scale. In this way, we destroy the process information storage only if is carried by the *j*^*th*^ scale, otherwise the information storage stays the same.

**Step 4:** With **Y**′ we compute again the *AIS*. We illustrated this step in Fig. 1, D. Let **Y**_*<t*_ be the set of past variables of the *process* previously found in the analysis, with 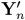 being the *n*-th *process surrogate* under analysis at scale *j*; then, the *AIS*′ for the surrogate data is:

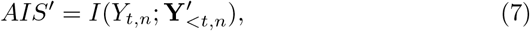

The algorithm is repeated from step 2 to step 4 for *n permutations*, with *n* = 1, …, *N*, to create a distribution of surrogate 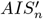 values; *N* is set according to the desired critical level for statistical significance (including Bonferroni correction for the number of scales, see below). Subsequently, all the *J*_0_ scales decomposed by the MODWT in step 1 are subjected to step 2, step 3 and step 4, such that *J*_0_ separate distributions of 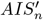 -values, one for each scale, are obtained.

**Step 5:** As a final step, the *AIS* is tested for statistical significance against the *J*_0_ different distributions of *AIS*′ surrogate values. If the **Y**^*j*^ (where *j* is one of the scales decomposed by the MODWT) contributes to the generation of the active information storage in the process **Y**, a significant drop of the *AIS*′ surrogates will be observed. This step is applied for all *J*_0_ scales under analysis and a Bonferroni correction is applied such that each individual scale is tested at the significance level *α/J*_0_.

Additionally, each scale analyzed is plotted, see Fig. 1, E, and we restrict ourselves to interpret only the scale that shows maximal distance (or well separated local maxima) from the original *AIS*, 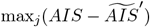, where 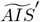 denotes the median of the surrogates distribution. We consider the maximal distance in addition to the statistical significance test because frequency decomposition is never perfect (e.g. due to leakage, noise and overlapping wavelet bands). Indeed, validation of the algorithm on synthetic data shows that the maximum distance reliably reflects the ground truth, whereas the statistical significance test can suffer from leakage effects on adjacent scales. Obviously, this limits the detectability of frequency-specific *AIS* and may be overly conservative. Thus, in scenarios, where AIS from multiple frequency bands is strongly expected *a priori*, or where the length of the data allows for vanishing leakage effects, the above restriction may be lifted.

## Results

In the following section we test the capability of the proposed algorithm to recover frequency specific AIS. To this end, we employed three simulations, where the ground truth is known. These simulations are limited to three example cases only, because the core idea and implementation strictly followed the spectral TE algorithm [10] (see above). For this more complex case of source-target interactions we have already demonstrated in depth that the MODWT construction of frequency specific surrogates in combination with a suitable statistical test reliably delivers a frequency resolved information theoretic measure [10].

In addition to the proof-of-principle on simulated systems, we applied the spectrally resolved AIS on Local Field Potential (LFP) data from ferrets under different levels of isoflurane and at recording sites in different cortical, and at sites in the prefrontal cortex (PFC) and in primary visual cortex (V1). For each combination of cortical area and layer we assessed if the AIS and the frequency resolved AIS were modulated as a function of different isoflurane concentrations using Bayesian linear regression.

All the analysis of AIS and spectrally resolved AIS below were performed with a block permutation of the wavelet coefficients (for construction of surrogates) and LA(8) as mother wavelet, similarly to [10].

### Example I: Null case, no information storage

At first, we simulated the case of no AIS in a process, to evaluate the behavior of our algorithm when none of the frequency scales generates information storage. We employed a white noise process, which by definition should not contain any information storage.

As expected, no significant AIS could be found in the time domain (see Fig 2, panel A). To evaluate the behavior of the spectral AIS algorithm in the scenario of no AIS we repeated the spectral analysis 500 times, to estimate the number of false positive result at each scale. This analysis revealed no false positive result, indicating that the strategy to exploit wavelet filtering for surrogates data creation leads to a conservative behaviour.

**Fig 2.**
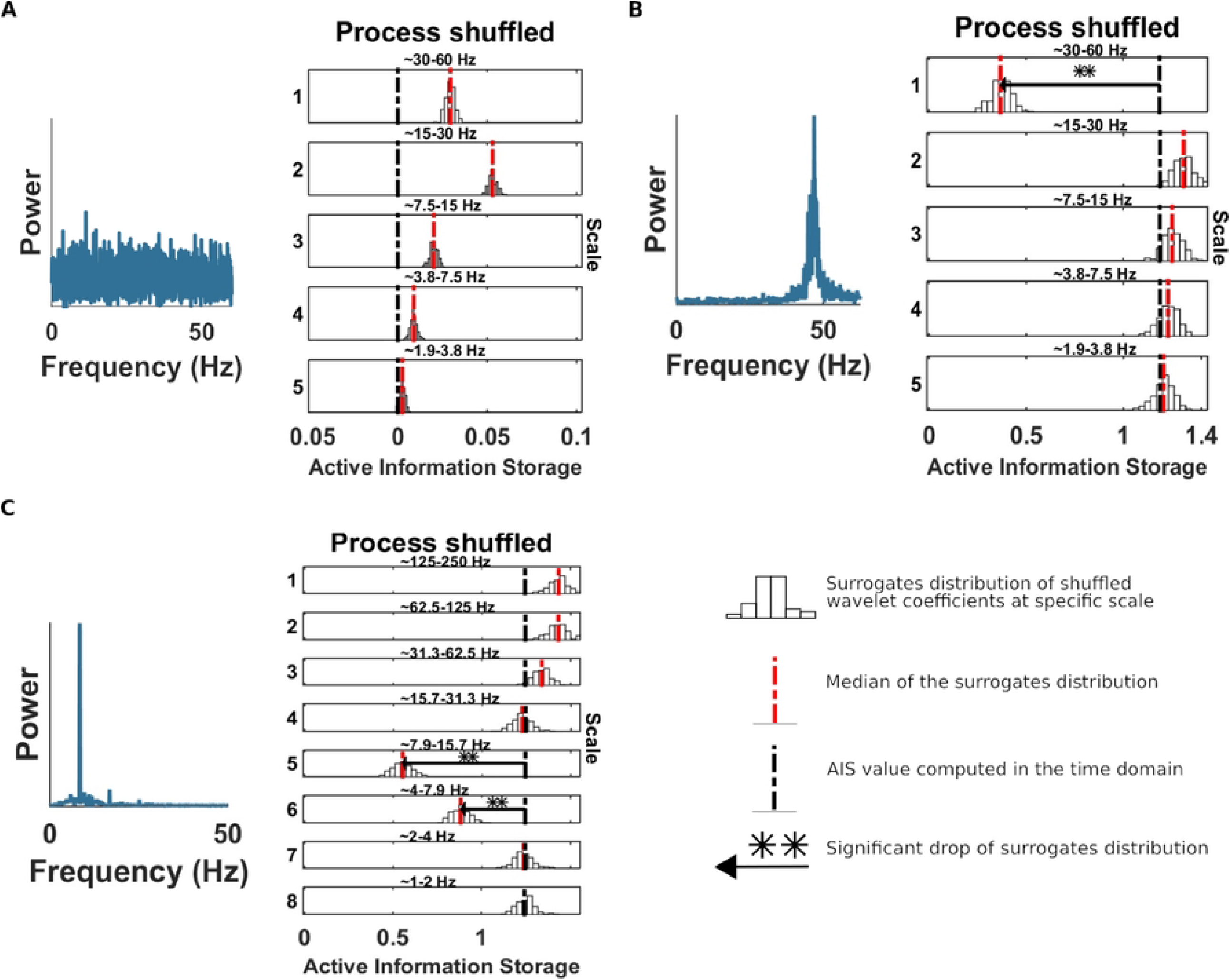
Spectrally-resolved AIS for three exemplary simulations. Each panel, shows the *AIS*′ distribution obtained from the surrogate data with shuffled coefficients at the scale indicated to the left, or, equivalently, the frequency band indicated at the top of each panel. White bars represent histograms of surrogate data, i.e. relative frequencies in (a.u.), the red dashed line is the median of the surrogate *AIS*′ distribution, the black dashed line is the original AIS value. The horizontal black line indicates the distance between the original AIS and the median of the surrogate distribution (**, *p <* 0.005; *, *p <* 0.05) Panel A, spectral AIS for the null-case example. Left, the power spectra of white noise process. Right, spectrally resolved AIS at different scales (frequency bands). No significant drop of the shuffled wavelet coefficients can be found since the process had no internal dynamic. Panel B spectral AIS for example 2, linear. Left, power spectra of AR(2) with spectral peak at 45 HZ. Right, spectrally resolved AIS at different scales (frequency bands). The AIS showed, correctly, a significant drop at scale 1 (30 −60Hz). Panel C, spectral AIS for example 3, nonlinear. Left, power spectra of the realizations of a selected variable of the Rossler system, with a spectral peak at around 8 Hz. Right, spectrally resolved AIS at different scales (frequency bands). The AIS showed, correctly, the largest drop at scale 5 (8 − 16 Hz) and scale 4 (4 − 8 Hz).

### Example II: AR process

Secondly, we simulated an autoregressive process AR(2) process with an autonomous oscillation at *f*_1_ = 45 Hz. We reproduced the simulation of Example 3 in [10], with *p* = 0.98, generating 10 s at sampling rate of 120 Hz and 10 trials, and closely following [34]. This process exhibits long-term memory and positive information storage [34]. The AR(2) process was generated with the following equation:

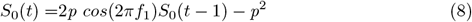

First, we analysed the process in the time domain to establish the presence of significant AIS. Then, we applied the spectrally-resolved AIS algorithm to obtain the frequency information of the system. Correctly, the scale 1 (frequency band 30–60 Hz), containing the spectral peak at 45 Hz, shows a significant drop of AIS in the surrogate data set indicating spectral AIS at that scale (see Fig 2, panel B).

### Example III: chaotic dynamical system oscillator

In this third simulation, we evaluated the spectral AIS with a process generated by a non-linear dynamical system that exhibits self-sustained periodic oscillations, similar to [10, 35]. The system was simulated with the following equation:

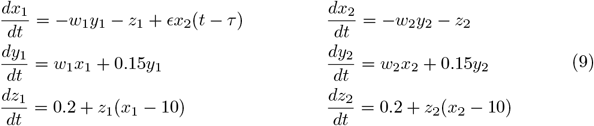

where *w*_1_ and *w*_2_ are the natural frequencies of the oscillator which were set to 0.8 and 0.9, and *ϵ* = 0.07 is the coupling strength and *τ* is the time delay, which was set to 2 time steps. Additionally, Gaussian white noise was added to the generated time-series. The analysis was performed on the assumption that only variables *x*_1_(*t*) could be observed. As can be seen in Fig 2, A, the process *x*_1_(*t*) oscillated around 8 Hz. The sampling rate was 500 Hz, and 25 trials of length 4 seconds were generated (100000 samples).

As before, first we established the presence of significant AIS in the time domain. Then, we obtained the spectral AIS as in the examples before. The results indicate that the largest drop was at scale 5, with also a significant drop at scale 6, which was expected as the frequency of the process spanned both scales (see Fig 2, panel C).

### Spectral AIS under anesthesia at different cortical layers via Bayesian regression

We applied our spectral AIS method to electrophysiological recordings of LFP data. Data were recorded in V1 and PFC in two different female ferrets, at supragranular layers, the granular layer and infragranular layers, under different concentrations (0.5%, 0.75%, 1%) of isoflurane and under awake conditions (0%).

These laminar LFP data have been analysed previously in terms of frequency spectrum modulations at different isoflurane concentration in [15]. Here, we provide a spectrally-resolved assessment of the AIS in these signals, we hypothesise that AIS is modulated by isoflurane concentration in a layer- and brain-region-specific way. All methodological and recording details can be found in [15].

### 0.0.1 AIS and spectral AIS estimation

First, we estimated the AIS from LFP recordings. We implemented a similar approach as in [8], using the IDTxl toolboox [21] to determine the presence of significant AIS value at the layer level. To make any claim about isoflurane concentration effects we had to use identical embeddings(see Section *Technical Background: Active information storage (AIS)*) for the estimation of the AIS or spectral AIS measures at different isoflurane levels in order to equilibrate the estimation bias across these levels. To this end, we applied the following four analysis steps:

1. Run the AIS algorithm for each trial (length 8 seconds) and isoflurane level.
2. Take the union of all embeddings across trials and isoflurane levels (i.e. the union of all past state variables identified).
3. Compute the AIS measure using the union embedding.
4. Apply the union embedding for the spectral AIS algorithm and quantify the frequency (scale/frequency band) contribution as:

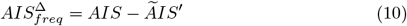

where, AIS is the original measure computed in the time domain and 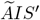 is the median of the AIS distribution estimated on surrogate data with coefficients shuffled at a specific frequency scale. Thus, the 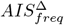 reflects the contribution of the particular frequency band under analysis to the AIS. Only positive values correspond to a significant contribution to the formation of AIS in the process under investigation.

### Bayesian Linear Regression layers: model specification

For the analysis of LFP laminar data, we employed a Bayesian linear regression model. The dependent variables were the *AIS* and *AIS*_*freq*_. For the *i* th trial, we can define the likelihood of the AIS measure as:

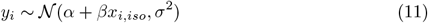

where *α* is the intercept and encodes the mean AIS, the parameter *β* is the slope which captures the isoflurane experimental effect, whereas the term 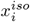 encodes the isoflurane levels (0%, 0.5%, 0.75%, 1%), and *σ*^2^ is the residual variance.

We choose a Normal distribution as a *prior* for the parameters *α* and *β* and Halfnormal distribution for the *σ* parameter, all values for the parameters of the *prior* distributions can be found in table S1. We built one model (we refer to this model as “simple model”) for each layer at PFC site and V1 site, separately (6 models in total). Since the plotted data showed a possible quadratic effect as a function of different isoflurane concentrations, we additionally built the same six models with an added quadratic term (we refer to this model as “model squared”), so that the likelihood terms become:

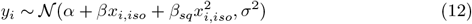

with a Normal distribution as a prior for *β*_*sq*_ (see table S1). Finally, for each layer we evaluated the model that predicted the data better (simple model vs model squared) using a leave-one-out cross-validation (LOO-CV) score as outlined next.

### Bayesian regression setup and model comparison

We estimated the model regression coefficients using Bayesian inference with Markov Chain Monte Carlo (MCMC) sampling, using the python package pymc3 [36] with NUTS (NO-U-Turn Sampling), using multiple independent Markov Chains. We implemented four chains with 3000 burn-in (tuning) steps using NUTS. Then, each chain performed 10000 steps, those steps were used to approximate the posterior distribution. To check the validity of the sampling, we verified that the R-hat statistic was below 1.05.

To evaluate different models with different numbers of parameters, we implemented cross-validation, which has been advocated for Bayesian model comparison, e.g. in [37]. In particular we adopted the LOO-CV implemented in PyMC3. Lower LOO-CV scores imply better models. We report the full modeling and model comparison results in 1 and only include the results of the winning models in the main text.

### Hierarchical Bayesian Regression

We point out that an alternative modeling approach to asses the anesthesia effects on the AIS measure would have been adopting a Hierarchical Bayesian Regression [38]. In a hierarchical model, parameters can be viewed as a sample from a population of parameters; for our case this implies to set a hyperprior from which we sample the *β* parameters for the two cortical areas (PFC and V1) and three layers (infragranular, granular and supragranular). This modeling approach would be optimal in case of the prior assumption of a certain amount of similarity of the AIS behaviour between these cortical structures, or in other words, an overall effect common to the different cortical areas and layers. The Bayesian framework allows to include such prior knowledge on the model formulation. However, previous work on spectral power [15, 16], as mentioned above, showed that anesthesia modulates cortical areas and layers differently. Based on this prior knowledge we decided to model each single layer in each cortical area (PFC and V1) separately, resulting in six separate models.

### Cortical layer and brain-region specific modulation of total AIS by isoflurane

We start by reporting the result of the Bayesian regression analysis for the *AIS* dependent variable, in the time domain, and subsequently the result of the frequency resolved AIS (*AIS*_*freq*_).

First, we evaluated the AIS measure for different isoflurane levels in the time domain, and performed the Bayesian regression analysis.

In V1, the models with a squared beta coefficient described the data better than the models without it, as indicated by the LOO-CV-based Bayesian model comparison **??** (lowest LOO score in supplementary Table S2). In contrast, in PFC, the two types of models were almost indistinguishable; yet the *model squared* performed slightly better as well (see supplementary Table S2).

In PFC, in the infragranular layer, we found a consistent increase of the AIS as a function of isoflurane concentration (yellow line) with a posterior mean of *beta iso* = 1.7, [1.28, 2.13] and *beta iso squared* = 0.9, [0.52, 1.27] (Fig 3, panel A) and also in the granular layer of PFC *beta iso*: = 0.48, [−0.075, 1.05] and *beta iso squared* = 1.43, [0.92, 1.91] (Fig 3, panel B), while for the super-granular layer of the PFC the effect of isoflurane on AIS was minimal, with a posterior mean of *beta iso* = 0.814, [0.08, 1.54] and *beta iso squared* = −0.54, [− 1.19, 0.099] (see Fig 3, panel C).

**Fig 3.**
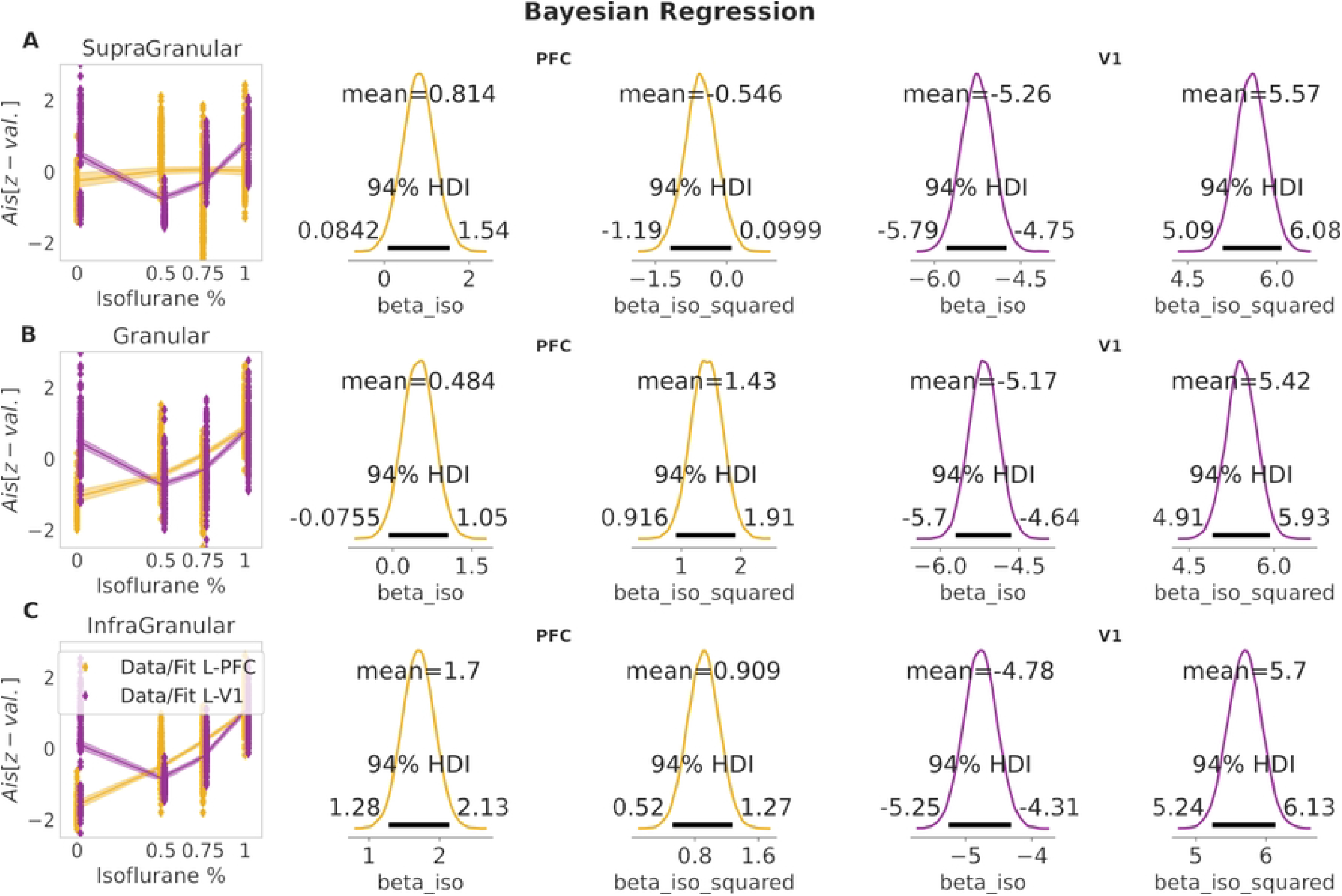
Bayesian Regression results of AIS in the time domain. Panel A, left, Bayesian regression fit for infragranular layer at PFC (yellow) and V1 (purple). Panel B, left, Bayesian regression fit for granular layer at PFC (yellow) and V1 (purple). Panel C, left, Bayesian regression fit for supragranular layer at PFC (yellow) and V1 (purple). Middle columns, posterior mean for *beta iso* and *beta iso squared* coefficients at PFC site, for panel A, B and C. Right columns, posterior mean for *beta iso* and *beta iso squared* coefficients at V1 site. Shaded area in the regression fit represents 94%HDI. Axis units are z-normalized values across anesthesia conditions.

In V1, all layers showed a similar behavior (see Fig 3, panels A–C), with a decrease of AIS values for intermediate isoflurane concentrations (0.5% and 0.75%) and a subsequent increase for the highest isoflurane level (1%). Posterior means for *beta iso* were =−4.78, [−5.24, −4.3], −5.17, [−5.69, −4.64] and −5.25, [−5.77, −4.73], for infragranular, granular and supragranular layers, respectively (see Fig 3, panel A–C). Similarly, posterior means for the beta coefficient of the squared isoflurane concentration were close to each other with *beta iso squared* = 5.7, [5.24, 6.13], 5.42, [4.91, 5.93], 5.57, [5.09, 6.08], for infragranular, granular and supragranular layers, respectively (see Fig 3, panel A–C).

In summary, deeper layers in PFC (infragranular and granular) showed stronger modulation under increasing isoflurane levels compared to supragranular layers, such a clear difference did not appear between layers in V1. This result is in line with [8], where a more pronounced increase of AIS (at increasing isoflurane concentrations) was found at PFC compared to V1.

### Cortical layer and brain-region specific modulation of frequency-specific AIS by isoflurane

Next, we evaluated the *AIS*_*freq*_ for different isoflurane levels in the frequency domain in multiple frequency bands. We start presenting the results for 62.5*Hz* − 125*Hz*, i.e. the high gamma frequency band.

In this band, in PFC, the *model squared* was substantially better than the *simple model* in the infragranular and granular layers; in the supragranular layer the *model squared*, despite still fitting the data better than the *simple model*, only had a marginally better LOO score (see supplementary Table S3).

In V1 at the infragranular layer, LOO scores for the models with or without the *beta iso squared* coefficient were almost identical, whereas for granular and supragranular layers the *model squared* represented a better description of the data by the model (see Table S3).

In the high gamma band (62.5*Hz* − 125*Hz*) we observed a modulation of the 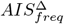 mostly in the supragranular layer of V1 (see Fig 4 panel A), with an increase for intermediate levels of isoflurane (0.5%) *beta iso* =−0.92, [−0.96, −0.89] and *beta iso squared* =− 0.86, [−0.89, −0.83] and a subsequent decrease for higher isoflurane concentrations, whereas in PFC such a modulation was absent in supragranular layer. Modulationwas was also absent at both, V1 and PFC, in granular and infragranular layers (see Fig 4, panels A–C).

**Fig 4.**
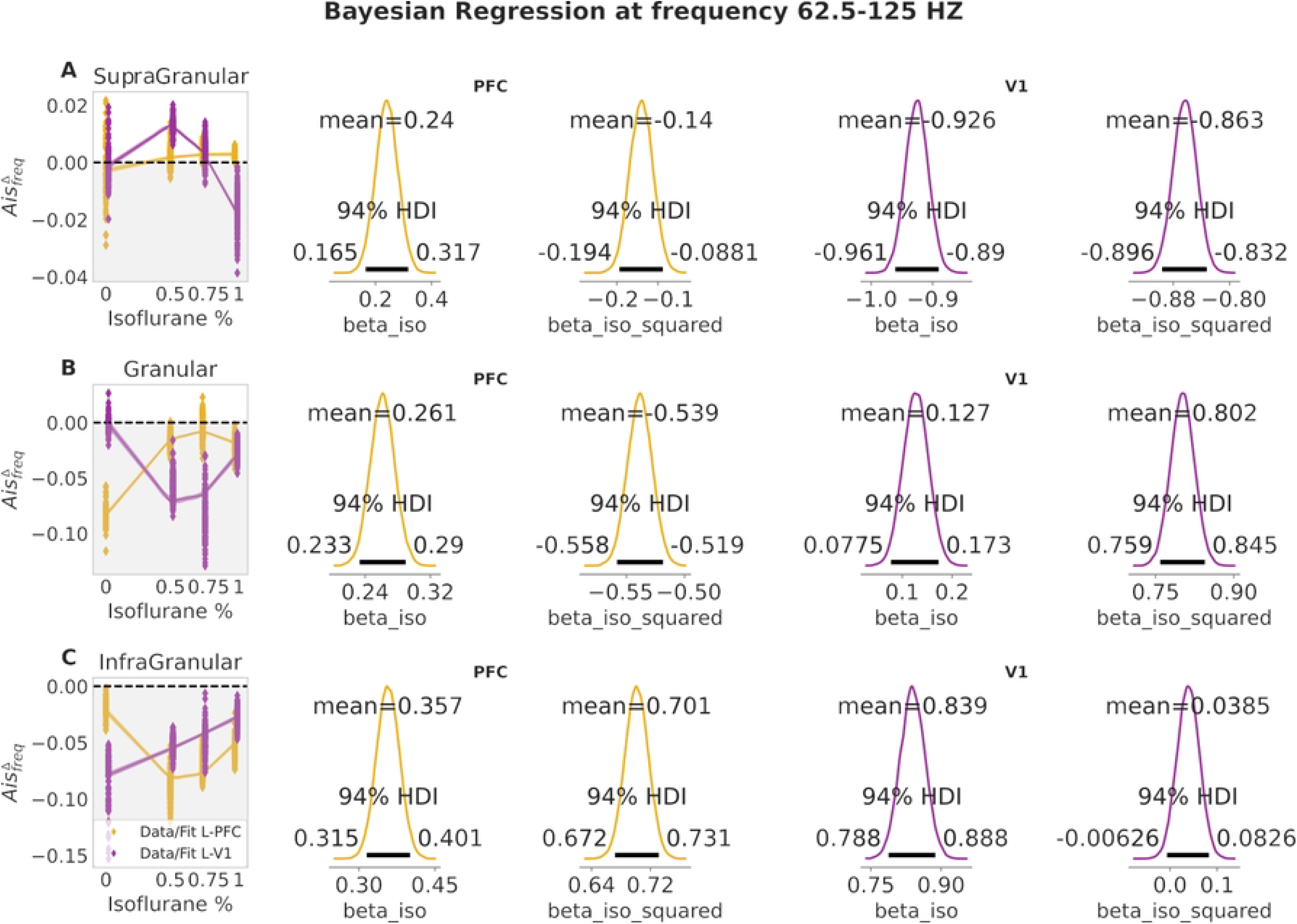
Bayesian Regression results of AIS at frequency 62.5-125 Hz. Panel A, left, bayesian regression fit for supragranular layer at PFC (yellow) and V1 (purple). Panel B, left, bayesian regression fit for granular layer at PFC (yellow) and V1 (purple). Panel C, left, bayesian regression fit for infragranular layer at PFC (yellow) and V1 (purple). Middle columns, posterior mean for *beta iso* and *beta iso squared* coefficients at PFC site, for panel A, B and C. Right columns, posterior mean for *beta iso* and *beta iso squared* coefficients at V1 site. Shaded area in the regression fit represents 94% HDI. Shaded gray background for 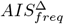 values that are below zero (i.e. no frequency specific drop)

In the frequency range 31*Hz* −62*Hz* (i.e. gamma band), the *model squared* was better for all layers at both brain regions (V1 and PFC). Nevertheless, at the granular layer of V1 and at the supragranular layer of PFC the LOO-CV difference with the *simple model* was minimal (see supplementary Table S4).

Similarly to the high gamma band, the supragranular layer was the layer most pronouncedly modulated by the different isoflurane concentrations. In the PFC the 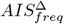 decreased as a function of isoflurane *beta iso* = −0.48, [−0.54, −0.41] and *beta iso squared* = −0.09, [−0.04, −0.14], while in V1 it increased for isoflurane at 0.5% followed by a decrease *beta iso* = −0.43, [−0.51, −0.35] and *beta iso squared* = −0.42, [−0.5, −0.36] (see Fig 5, panel A).

**Fig 5.**
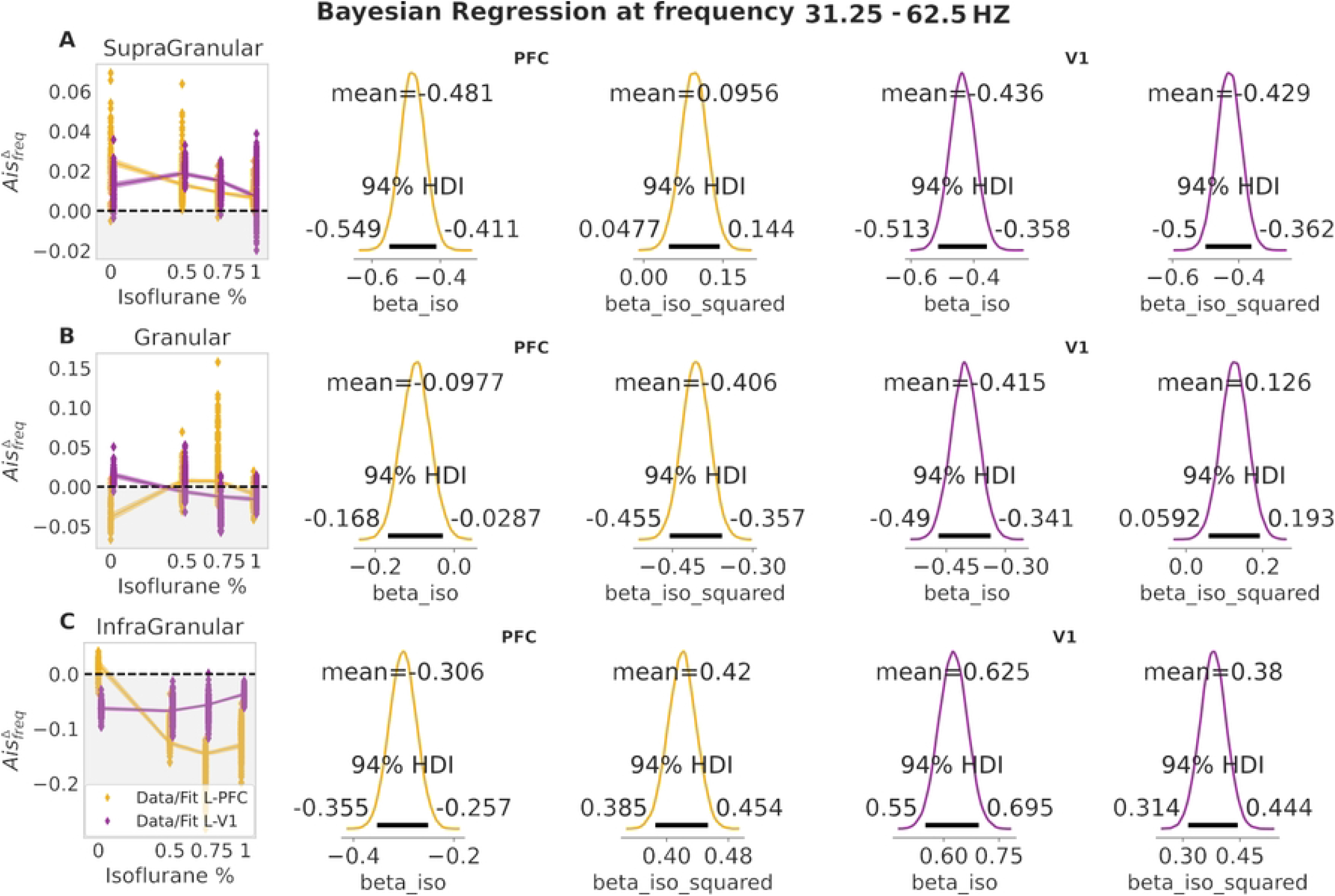
Bayesian Regression results of AIS at frequency 31.25-62.5 Hz. Panel A, left, bayesian regression fit for supragranular layer at PFC (yellow) and V1 (purple). Panel B, left, bayesian regression fit for granular layer at PFC (yellow) and V1 (purple). Panel C, left, bayesian regression fit for infragranular layer at PFC (yellow) and V1 (purple). Middle columns, posterior mean for *beta iso* and *beta iso squared* coefficients at PFC site, for panel A, B and C. Right columns, posterior mean for *beta iso* and *beta iso squared* coefficients at V1 site. Shaded area in the regression fit represents 94% HDI. Shaded gray background for 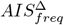 values that are below zero (i.e. no frequency specific drop)

At granular layer of V1, the positive 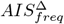 values for isoflurane at 0% decrease to negative (see Fig 5, panel B, shaded gray background) for intermediate and high level of isoflurane concentrations (from 0.5% to 1%).

Finally, no relevant modulation could be seen in the infragranular layers of both PFC and V1 brain areas (see Fig 5, panel C). Taken together, the results for gamma and high gamma band, showed an isoflurane effect on 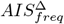 mainly in the superficial layer (supragranular) and minimally in the granular layer, in agreement with association of gamma band to superficial layers [**?**], see Section *Modulation of spectral information storage according to distinct functional roles across cortical layers by anesthesia*, for further details.

In the frequency range 15*Hz* −31*Hz* (i.e. beta band), the *simple model* had a lower LOO-CV score for the supragranular layer of PFC. In all other cases, the *model squared* had lower LOO-CV score (see Table S4).

In this frequency band, in PFC, the supragranular and infragranular layers decreased as isoflurane levels increased. While in the supragranular layer 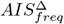 value were still positive at isoflurane 1%, *beta iso* = −0.67, [−0.72, −0.62], in the infragranular layer the 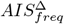 values became negative at higher isoflurane levels (0.75% and 1%),*beta iso* = −0.74, [− 0.79, −0.70] and *beta iso squared* = 0.11, [0.08, 0.16], revealing that the spectral surrogate drop was abolished (see Fig 6, panel C, shaded gray background).

**Fig 6.**
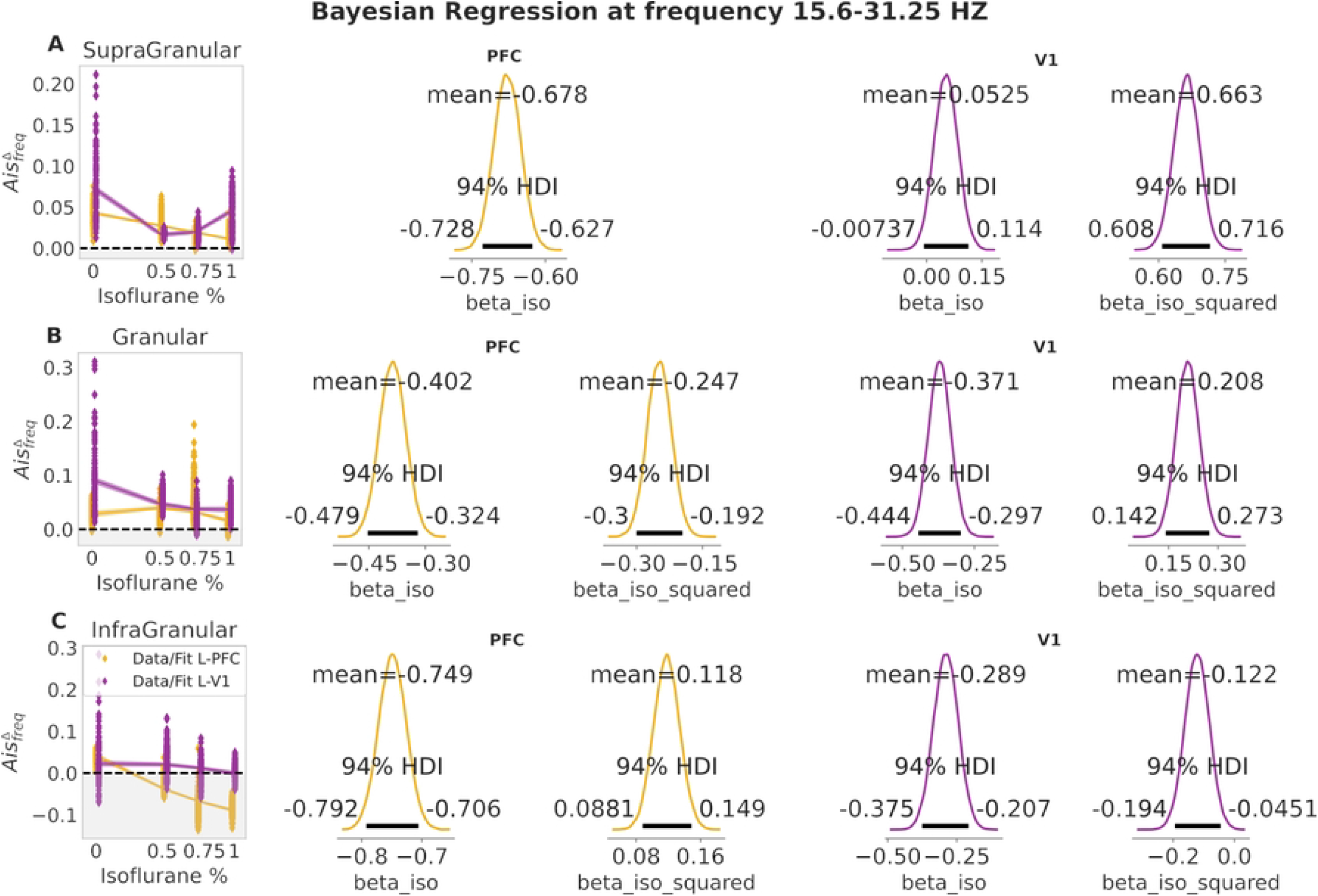
Bayesian Regression results of AIS at frequency 15.6-31.25 Hz. Panel A, left, bayesian regression fit for supragranular layer at PFC (yellow) and V1 (purple). Panel B, left, bayesian regression fit for granular layer at PFC (yellow) and V1 (purple). Panel C, left, bayesian regression fit for infragranular layer at PFC (yellow) and V1 (purple). Middle columns, posterior mean for *beta iso* and *beta iso squared* coefficients at PFC site, for panel A, B and C. Right columns, posterior mean for *beta iso* and *beta iso squared* coefficients at V1 site. Shaded area in the regression fit represents 94% HDI. Shaded gray background for 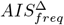 values that are below zero (i.e. no frequency specific drop)

In V1 the supragranular layer had a different modulation, compared to PFC, with a significant decrease at isoflurane 0.5% and a subsequent increase for isoflurane 0.75% and 1%,*beta iso* = 0.05, [−0.007, 0.11] and *beta iso squared* = 0.66, [0.60, 0.71] (see Fig 6, panel A). In granular layer after a decrease for concentration 0.5%, 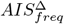 remained in a similar range of values for high isoflurane concentrations (see Fig 6, panel B).

In the frequency range 8*Hz* − 16*Hz* (alpha/beta band), the *model squared* had a lower LOO-CV for all layers at both PFC and V1 regions (see Table S5). Nevertheless, for the supragranular layer at PFC and granular layer of V1 the models had overlapping standard errors of the estimates (SE).

In this frequency range, in V1, we observed a similar behavior of supragranular and granular layers to the previous frequency range, posteriors for *beta iso* =−0.06, [−0.008, −0.125] and *beta iso squared* = 0.687, [0.63, 0.73] of supragranular layers were very close to the ones in the frequency range 15*Hz*−31*Hz* (compare values with Fig 7, panel A). In contrast, the infragranular layer of V1 had a opposite modulation to that found in the next higher frequency range with an increase for intermediate isoflurane level (0.5%) and a later decrease at higher isoflurane concentrations, *beta iso* =−0.574, [−0.64, −0.50] and *beta iso squared* = 0.52, [0.588, 0.46] .

**Fig 7.**
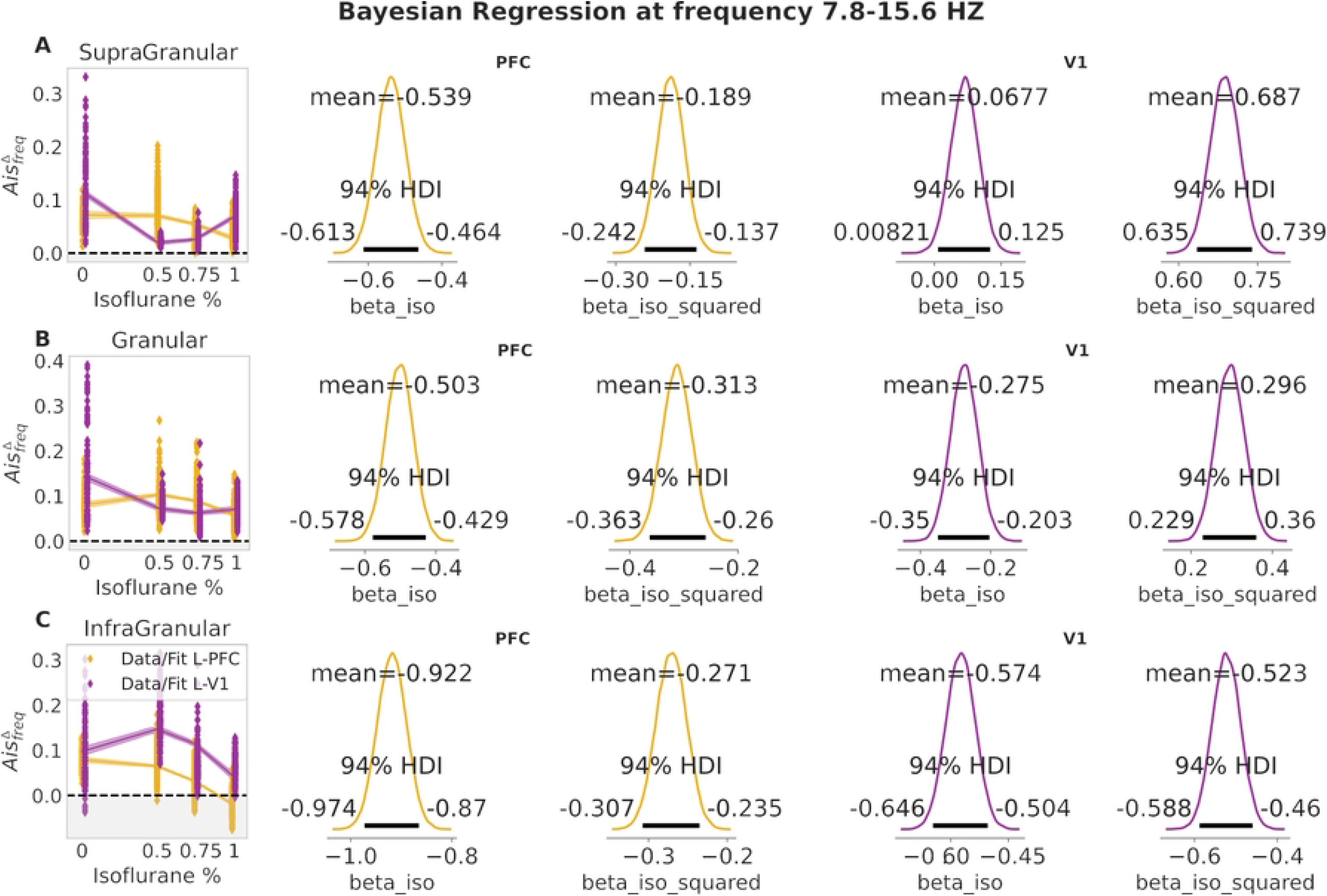
Bayesian Regression results of AIS at frequency 7.8-15.6 Hz. Panel A, left, bayesian regression fit for supragranular layer at PFC (yellow) and V1 (purple). Panel B, left, bayesian regression fit for granular layer at PFC (yellow) and V1 (purple). Panel C, left, bayesian regression fit for infragranular layer at PFC (yellow) and V1 (purple). Middle columns, posterior mean for *beta iso* and *beta iso squared* coefficients at PFC site, for panel A, B and C. Right columns, posterior mean for *beta iso* and *beta iso squared* coefficients at V1 site. Shaded area in the regression fit represents 94% HDI. Shaded gray background for 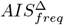 values that are below zero (i.e. no frequency specific drop)

In PFC the 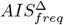 was mainly modulated from 0.5% to 1% isoflurane levels and this modulation was strongest in the infragranular layer (compare Fig 7, panel A, B and C yellow). Despite a common shift of alpha power from posterior to anterior cortex during loss of consciousness (LOC) [18], none of the layers at PFC showed an 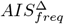 component increase. We discuss the absence of the alpha anteriorization effect commonly observed during LOC in Section *Modulation of spectral information storage according to distinct functional roles across cortical layers by anesthesia*.

In the frequency range 4*Hz* −8*Hz* (theta band), the *model squared* was substantially better only in the supragranular layer of V1 (see Table S6); in all other cases the difference with the *simple model* was minimal, yet the *model squared* had a nominally lower LOO-CV score.

In PFC, we observed a modulation of the 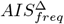 by isoflurane, in all three layers (see Fig 8, panel A-C). The strongest decrease was for infragranular and supragranular layers with *beta iso* =−0.515, [0.57, −0.45] and *beta iso squared* = 0.18, [0.14, 0.22] and *beta iso* =−0.301, [−0.375, −0.228] and *beta iso squared* = 0.151, [0.09, 0.203], respectively.

In V1 the supragranular layer had a strong decrease for isoflurane 0.5% but this was followed by an increase for higher isoflurane levels *beta iso* = 0.618, [0.56, 0.67] and *beta iso squared* = 0.776, [0.725, 0.827].

**Fig 8.**
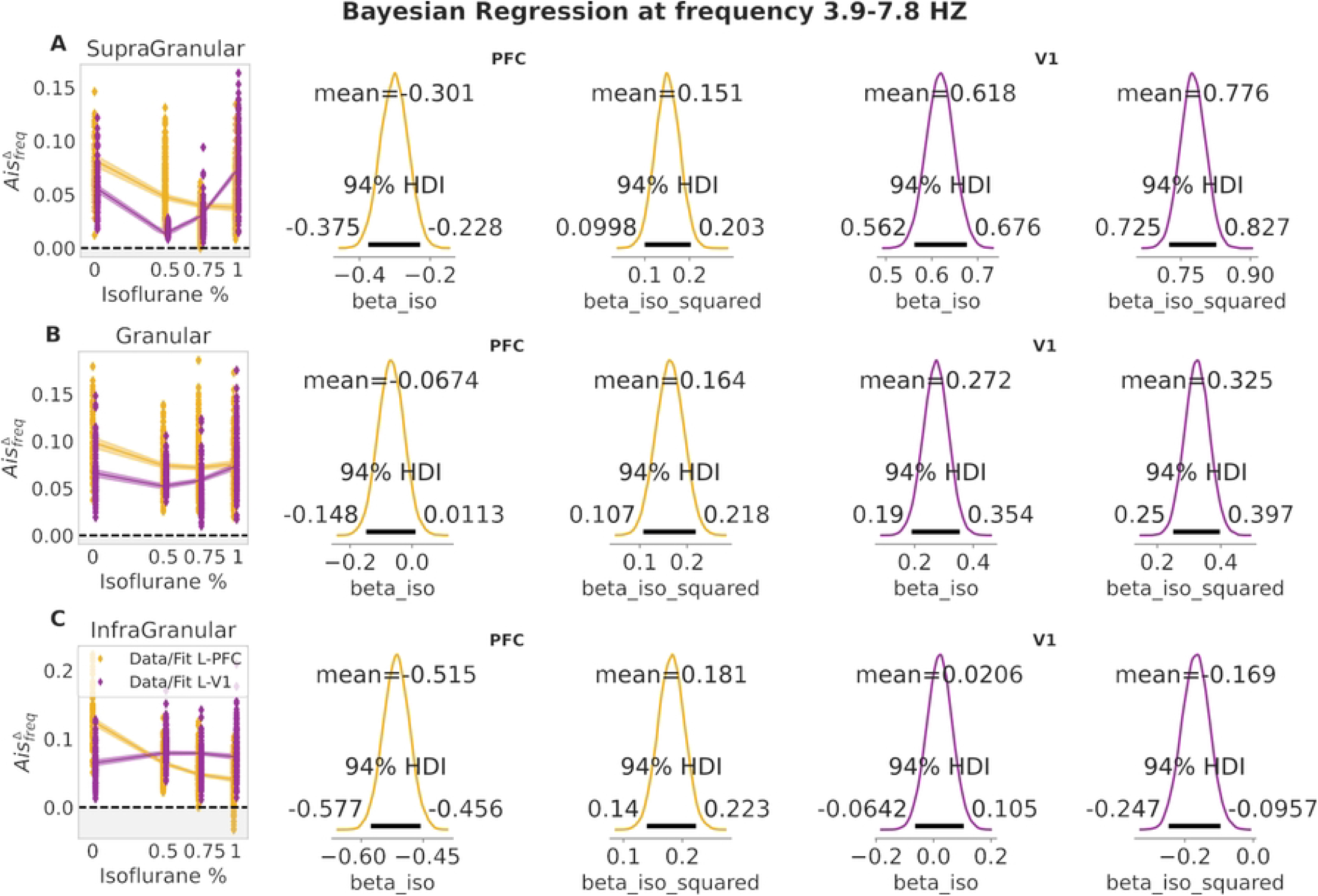
Bayesian Regression results of AIS at frequency 3.9-7.8 Hz. Panel A, left, bayesian regression fit for supragranular layer at PFC (yellow) and V1 (purple). Panel B, left, bayesian regression fit for granular layer at PFC (yellow) and V1 (purple). Panel C, left, bayesian regression fit for infragranular layer at PFC (yellow) and V1 (purple). Middle columns, posterior mean for *beta iso* and *beta iso squared* coefficients at PFC site, for panel A, B and C. Right columns, posterior mean for *beta iso* and *beta iso squared* coefficients at V1 site. Shaded area in the regression fit represents 94% HDI. Shaded gray background for 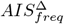 values that are below zero (i.e. no frequency specific drop)

In the frequency range 1.95*Hz* − 4*Hz* (delta band), the models with a squared beta coefficient described the data better than the models without it in all layers at PFC and V1, as indicated by LOO cross-validation-based Bayesian model comparison (lowest LOO-CV score, see Table S7).

At this low frequency band the modulation by isoflurane became more homogeneous across layers and brain regions. Indeed, we found that the infragranular and granular layer of PFC and V1 were all similarly modulated (see posterior plots in Fig 9, panel B and C). Only in supragranular layers the increase for high isoflurane values was stronger in V1 compared to PFC, with *beta iso* = 0.609, [0.55, −0.66] and *beta iso squared* = 0.77, [0.72, 0.82] for V1 and *beta iso* = 0.297, [0.22, 0.37] and *beta iso squared* = 0.391, [0.33, 0.44].

**Fig 9.**
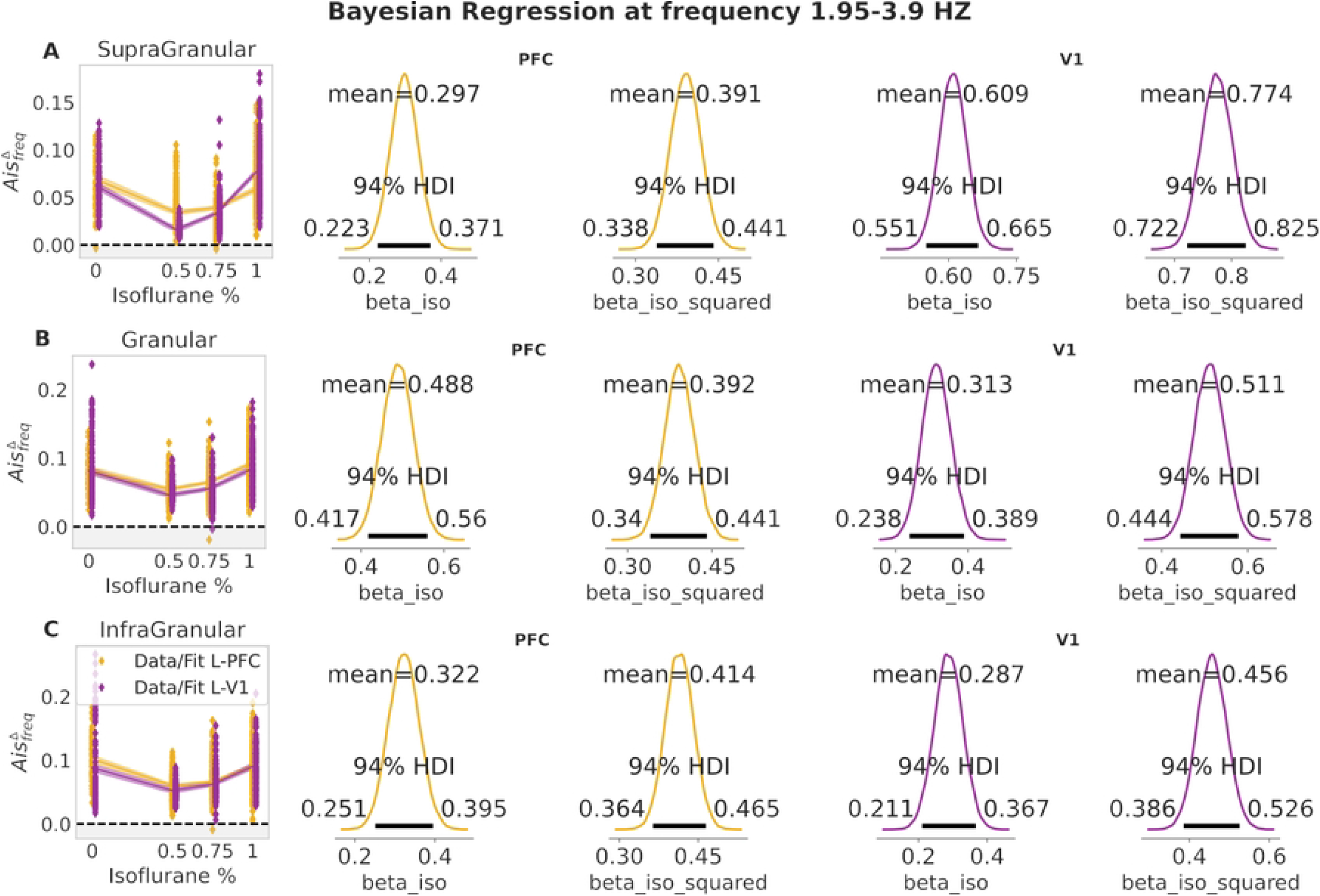
Bayesian Regression results of AIS at frequency 1.95-3.9 Hz. Panel A, left, bayesian regression fit for supragranular layer at PFC (yellow) and V1 (purple). Panel B, left, bayesian regression fit for granular layer at PFC (yellow) and V1 (purple). Panel C, left, bayesian regression fit for infragranular layer at PFC (yellow) and V1 (purple). Middle columns, posterior mean for *beta iso* and *beta iso squared* coefficients at PFC site, for panel A, B and C. Right columns, posterior mean for *beta iso* and *beta iso squared* coefficients at V1 site. Shaded area in the regression fit represents 94% HDI. Shaded gray background for 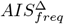 values that are below zero (i.e. no frequency specific drop)

Finally, we estimated the frequency range 0.9*Hz*−1.9*Hz* (low delta band). The Bayesian model comparison revealed that the *model squared* had a lower LOO score in all the layers, however in the granular layer of V1 standard error of estimates overlapped, indicating that both models fitted the data similarly (see Table S9).

As in the previous frequency range, deep and superficial layers at both brain regions were characterized by a similar isoflurane modulation, due to a possible global effect of slow oscillation throughout the cortex under LOC (see Section *Modulation of spectral information storage according to distinct functional roles across cortical layers by anesthesia*). However, in this frequency range, infragranular and granular layers at PFC had a stronger increase at high isoflurane concentrations (0.75% and 1%) than in V1, with the highest difference in the infragranular layer *beta iso* = 0.84, [0.78, 0.903] and *beta iso squared* = 0.284, [0.24, 0.32] for PFC and *beta iso* = 0.291, [0.20, 0.37] and *beta iso squared* = 0.25, [0.18, 0.33], while supragranular layer showed a similar modulation at both brain regions (see Fig 10, panel A).

**Fig 10.**
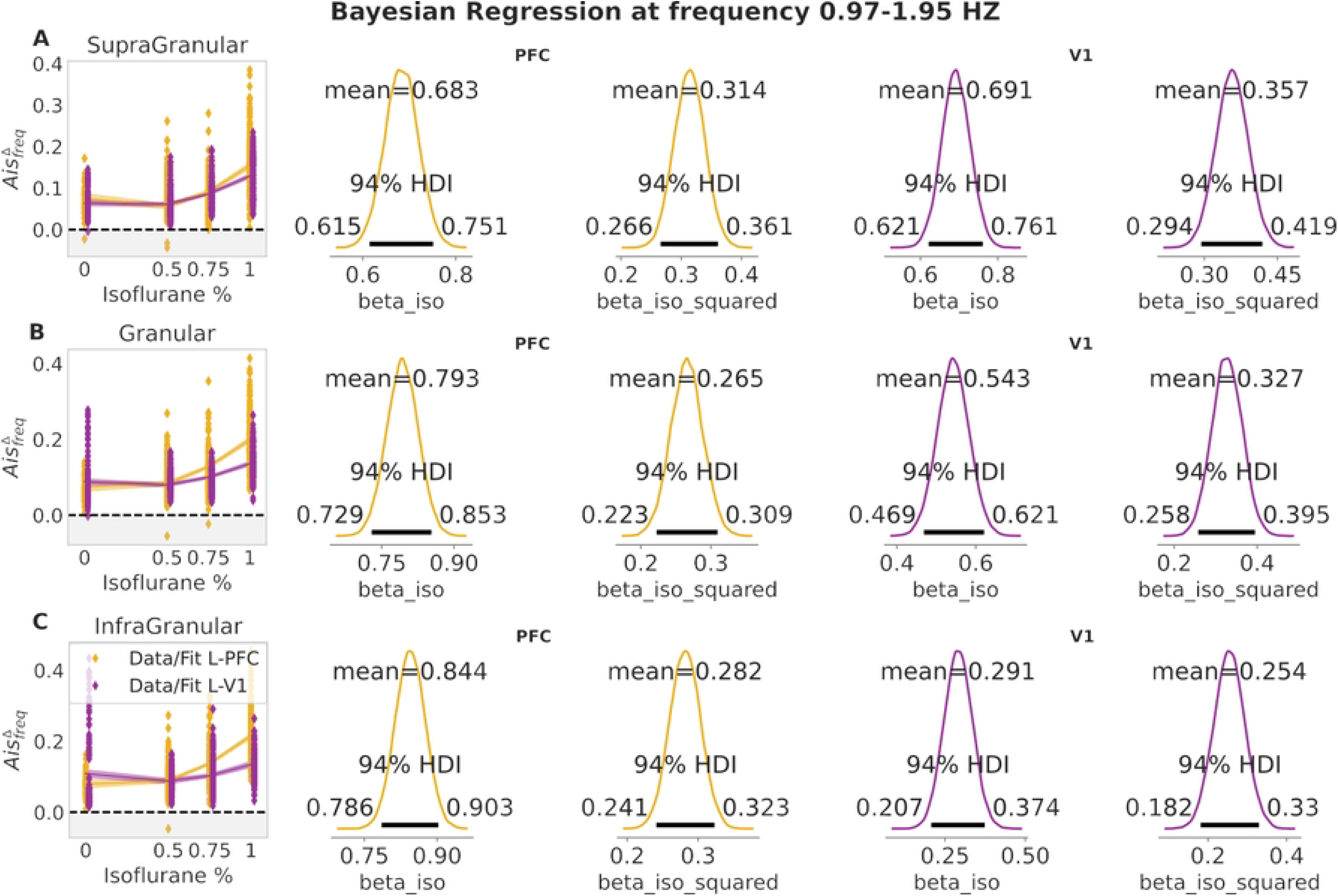
Bayesian Regression results of AIS at frequency 0.97-1.95 Hz. Panel A, left, bayesian regression fit for supragranular layer at PFC (yellow) and V1 (purple). Panel B, left, bayesian regression fit for granular layer at PFC (yellow) and V1 (purple). Panel C, left, bayesian regression fit for infragranular layer at PFC (yellow) and V1 (purple). Middle columns, posterior mean for *beta iso* and *beta iso squared* coefficients at PFC site, for panel A, B and C. Right columns, posterior mean for *beta iso* and *beta iso squared* coefficients at V1 site. Shaded area in the regression fit represents 94% HDI. Shaded gray background for 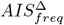values that are below zero (i.e. no frequency specific drop)

### 0.0.2 Relation between the spectral AIS and the spectral power

To further highlight the specificity of the spectral AIS compared with a simple spectral power analysis we performed a Bayesian correlation of two frequencies range: 0.9*Hz*−1.9*Hz* (delta) and 7.9*Hz*−15*Hz* (alpha). The Bayesian correlation revealed that the correlation between the two measures varied depending on the frequency bands or isoflurane level. In the frequency range 0.9*Hz* 1.9*Hz* at isoflurane of 0% there was a strong evidence for a negative correlation BF= 48530.5 (see Fig 11, panel A, top row); at isoflurane of 1% the correlation dropped to moderate evidence BF= 3.079 (see Fig 11, panel A, bottom row). In the alpha band 7.9*Hz*−15*Hz* at isoflurane of 0% the evidence for a correlation was absent BF= 0.098 (see Fig 11, panel B, top row) and became positive with weak or anecdotal evidence BF= 1.380 (see Fig 11, panel B, bottom row). Additionally, we showed that as the power increase as a function of isoflurane the 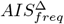 had opposite behavior; the frequency specific surrogates drop increased at higher isoflurane percentage in the range 0.9*Hz* − 1.9*Hz*, while it decreased in the range 7.9*Hz* − 15*Hz* (see 11, panel C and D).

**Fig 11.**
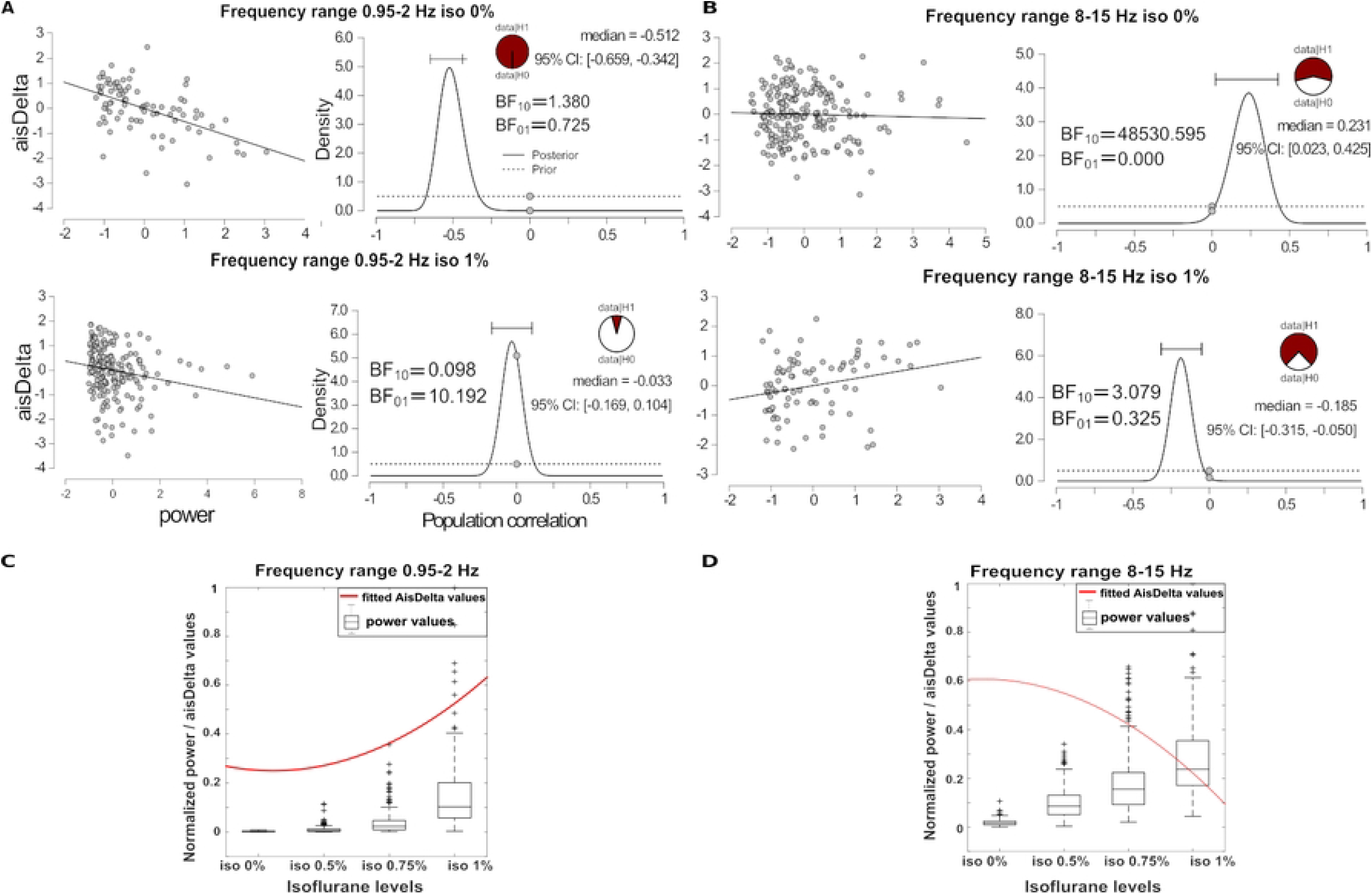
Bayesian correlation of spectral AIS and spectral power. Panel A, bayesian correlation of spectral AIS with spectral power in range 0.9*Hz*−1.9*Hz* at isoflurane 0%, top row and isoflurane 1% bottom row. Panel B, bayesian correlation of spectral AIS with spectral power in range 7.9*Hz*−15*Hz* at isoflurane 0%, top row and isoflurane 1%, bottom row. In each panels A and B, on the right: correlation plot of 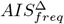 values, on the left: posterior distribution (black line), prior distribution (dashed line), median and 95% of the posterior estimates and bayes factor BF10 of the alternative hypothesis (H1) and BF01 of the null hypothesis (H0). Panel C, the modulation of AIS with isoflurane levels (red curve) follows the increase of spectral power in the delta band (black box-plot). Panel D, the modulation of AIS with isoflurane levels has an opposite behaviour (red curve) with the modulation of spectral power in the alpha band (black box-plot).

## Discussion

In this study we addressed two tightly related questions: 1. Can we in principle design an algorithm to detect frequency-specific active information storage? 2. Do we detect such frequency-specific active information storage in neural systems, and if so, does it provide valuable information towards a better understanding of neural information processing.

### Estimating spectrally-specific AIS

To address the first question, we presented an algorithm to estimate which specific spectral components contribute to the overall active information storage (AIS) of a process. We demonstrated with different simulations that these spectral components can be reliably identified, in both linear and nonlinear systems and processes. The algorithm builds on the idea of creating spectrally specific realizations of the null model (surogate data) that was presented in [10], for the case of spectrally-resolved TE. In the present study, we again used the MODWT decomposition and scrambling of the wavelet coefficients for the creation of spectrally-specific surrogates data to asses frequency contribution to the AIS measure.

The spectrally-resolved AIS can be seen as an attempt to determine frequency components related to information-theoretic properties of a process under investigation. Its estimation is less complex than the estimation of spectral TE which implies to decompose specific spectral components of sources and targets and to distinguish one-to-one from one-to-many or many-to-one interactions, whereas spectral AIS provides spectral resolution only of a single process. Thus, it is expected that the proposed algorithm performs similarly well as the previously published one for spectral TE. We, therefore, keep the discussion of the validity and performance short here.

### Which insights into neurophysiology can be obtained using spectral AIS? – The example of anesthesia effects

The second question becomes crucial in biological systems such as the brain where the role of rhythmic processing in neural system is still not fully understood but prominently discussed for neural communication mechanisms [39–41]. We note that for AIS as a foundational component of neural information processing, only a handful of studies exist to date [**?**, 6–8, 13], and none has asked the question of its relation to neural rhythmic processing.

The analysis of LFP cortical layers at two brain sites (PFC and V1) in ferrets, showed that we can successfully detect frequency-specific AIS at different frequency bands. While in the time domain the total AIS showed an increase as a function of isoflurane level, the spectral perspective revealed a much more complex and richer picture. Furthermore, in comparing modulations of spectral AIS and spectral power we showed that AIS provides information on the computational dynamics of the neural process and its modulation by anesthesia, which spectral power analysis does not directly reveal.

In the remainder of this section, we will further discuss the results of the application of our novel, spectrally-resolved AIS measure to the LFP, highlight the additional details that a spectral decomposition revealed compared to time-domain-only analysis and the additional information that the spectral AIS provides compared to only spectral power analysis also in terms of anesthesia effects. We will then indicate limitations and caveats of the method, discuss its relation to previous approaches and possible future applications.

### Modulation of spectral information storage according to distinct functional roles across cortical layers by anesthesia

Even though the molecular targets of anesthetic agents are well known [42], changes in neural information processing related to anesthesia are not well understood. This understanding is further complicated by the complex laminar organization of the cortex, which is composed of neurons with distinct connectivity patterns and feedforward or feedback projections across laminae [16, 17, 43–45]. Of importance to our study is also the fact that neural assemblies and canonical circuit motifs exhibit rhythmic activity in distinct frequency bands [40] where activity supragranular layers is found to be often dominated by gamma band activity and projected mostly along the feedforward direction of the cortical hierarchy, while activity in infragranular layers is often dominated by alpha/beta frequencies and projected along the feedback direction [43]. Note however, that recent experiments using different stimuli challenge this clear link of frequency bands to layers [45]. Thus, information theoretic quantities in cortex and their anesthesia related changes are bound to exhibit a complex spectral structure – possibly in a layer-specific way.

Our results indicate that also the spectral AIS reflects the previously described frequency signatures within a cortical area. Modulation by isoflurane could be seen in the high gamma range (62.5*Hz*− 125*Hz*) in superficial layers (e.g. supragranular) of V1 and in the gamma range (31*Hz*− 62.5*Hz*) at both V1 and PFC areas. In beta and alpha frequency ranges the strongest modulation effect were observed mainly at the PFC area in deeper layers (e.g. granular and infragranular). Departing from the simple, previously reported, associations between layers and frequencies the supragranular layer of V1 showed a decrease in AIS at intermediate isoflurane concentration, followed by an increase at higher concentrations in a broad frequency range from 4 Hz to 30 Hz. This means that in supragranular layers, also information processing in non-gamma frequencies is affected by anesthesia. This pattern specifically was actually more pronounced in the supragranular that in granular and infragranular layers. In contrast to these region and layer specific anesthesia effects at frequencies above 4Hz, at ferquencies below 4 *Hz*, the differences between brain regions (PFC or V1) and layers became minimal. This could mean that the emergence of slow-wave oscillatory power has to have an indiscriminate global effect on the neural computational properties throughout the cortex. In the case of slow oscillations below 2 *Hz*, the PFC showed a similar pattern at all three layers with a substantial increase of the 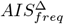 towards more positive values at the high isoflurane level (1%), with the strongest increase at deep infragranular layers in agreement with [46]. Our results are in line with previous works reporting that with LOC, neural signals became more uniform and exhibited repetitive patterns, interrupted by bursting activity [47]. In particular, slow-wave oscillatory power was more pronounced during anesthesia-induced LOC.

A potential explanation for the observed differences in isoflurane modulations of local information processing at the two cortical areas, V1 and PFC, may be the differences in the quantitative properties of neural circuits in the two areas. The different properties of individual neurons, feedback loops and feedforward excitation or inhibition from other areas (e.g. thalamus) might affect how information is stored and maintained at the local level under different anesthesia concentrations. Additionally, the functional organization of these areas might play a role, for example missing sensory input during LOC at V1 or reduced integration in PFC that serves higher cognitive function [48]. Studies that investigate neural processing in animals under anesthesia will have to consider that different cortical sites can be modulated or affected by anesthesia differently, and that the local processing exhibits distinctive spectral AIS, which in turn might affect how information is integrated and transferred across brain regions, as previously also reported by Wollstadt et al. [8].

### Spectrally-resolved AIS provides insights into neural processing and the effects of anesthesia that are not provided by an analysis of spectral power

Anaesthetic agents such as isoflurane, sevoflurane or propofol produce similar oscillatory changes, in particular, predominating low frequencies (delta band) and increased power in the alpha frequency band at frontal sites (anteriorization effect) [18, 49, 50]. We show here that the spectral AIS provides additional information on the underlying neural information processing and the effects of isoflurane. For example, we describe effects in the alpha frequency band that are not found by an analysis of only spectral power: While at low frequencies (delta band) and no isoflurane (0%), the spectral AIS and the spectral power are correlated, such a relation can not be seen in the alpha band (Figure 11 A, B). Additionally, while in the delta band the spectral AIS follows the increase of spectral power as a function of isoflurane concentration, an opposite behaviour can be seen in the alpha band (Figure 11 C, D).

Thus, decomposing the AIS measure in its spectral components can reveal aspects of the computational dynamics of neural processes that are not directly accessible by a spectral power analysis. This will be highly useful for anesthesia research in general, e.g., also for other anesthetics such as propofol. For propofol, recent work showed that alpha band effects (increase of anterior alpha and decrease of posterior alpha) depended on two types of thalamocortical circuits affected by the anaesthetic agents and were completely distinct from the propofol-induced slow oscillations [50]. Thus, we speculate that also propofol-related changes in spectral AIS will be distinct for the delta- and the alpha-bands.

### Spectrally-resolved AIS adds additional insights into the effects of anesthesia on neural processing compared to AIS in the time domain

In a previous study analyzing LFPs from ferrets under anesthesia, the AIS (in the time domain) increased as a function of isoflurane concentrations in PFC [8]. Given the similar effect that we found in this work in the time domain (see Fig 3, panel C), the overall increase at high isoflurane levels in AIS seems to be linked to an increase in AIS in delta frequencies, whereas alpha and beta frequency bands are modulated by isoflurane differently, i.e. they decrease as a function of isoflurane levels. Alpha and beta bands have been linked to generation of internal models in the predictive coding framework [13], and have also been associated mostly with deep cortical layers, and thereby, cortical feedback pathways [16]. Thus, the absence of alpha and beta-band AIS may suggest—following the line of argument in [13]—that under anesthesia the maintenance of internal models and the generation of internal predictions is strongly impaired. This in turn may be an important component of the phenomenon of loosing consciousness.

### Relation to previous approaches

A method that allows to compute AIS at different temporal scales has been introduced by [51]. It exploits the state space formalism to obtain a multiscale representation of a linear fractionally integrated autoregressive process (ARFI) [51]. The time-series undergo to lowpass filtering and downsampling to obtain a multiscale representation, so that the AIS can be computed as a function of the cutoff frequency. This parametric formulation, employing the state space formalism is restricted to the description of linear Gaussian processes. However, it is a significant improvement over previous attempts to quantify system complexity in terms of a linear multiscale entropy [52], with the simultaneous description of short and long memory properties which are fundamental aspects of systems dynamic [34]. When the assumptions of linear Gaussian processes is valid the approach from [51] will be more data-efficient and come with lower computational burden.

### On the possibility of cross-spectral information storage

Due to the sensitivity of information-theoretic measures to non-linear phenomena it is conceivable to find information storage in cases where the frequency of the process underlying the storage changes over time, i.e. where the stored information wanders between frequencies as the process unfolds. If, for example, the information is moving forth and back between dynamics at certain low frequencies and certain high frequencies, this should be detectable. Thus, as an extension to the algorithm presented here, it is possible to destroy the information in a specific frequency also in the future of a process instead of the past, similar to the individual frequency-specific destruction of information in source and target processes in the estimation of TE [10]. Although this is not used in the current manuscript, it is implemented in *IDT*^*xl*^ [21], for future investigations.

### Caveats and limitations

The estimation of information-theoretic quantities, such as the AIS, from finite data is highly non-trivial (e.g. [53] and references therein). In many cases the necessary number of physical realizations of a process is not available. Two possible strategies can be implemented then: pooling data over time to obtain a sufficient amount of realizations (this requires stationarity) or pooling data over an ensemble of temporal copies. This latter approach approach exploits the cyclostationarity across these temporal replications of the process. Last, for discrete-valued data, Bayesian approaches exist for optimization embedding parameters and AIS estimation [54]; these approaches are available in our Toolbox [21].

### Future directions

Future studies should focus on combining spectrally resolved transfer entropy [10] and active information storage to provide a more exhaustive characterization on the computational behaviour of the analysed system in the spectral domain. Employed together, these tools offer a promising framework to test specific hypothesis on brain functioning such as predictive coding theory [9] or encoding and maintenance of information in working memory [55]. For example, frequency-resolved measures of information transfer and active information storage can test specific hypothesis on LFP-frequency signatures of error signals [17, 43] or coding of prior information [13]. Similarly, maintenance of relevant information for later reactivation, in working-memory and prefrontal cortex has been associated with specific frequency signature [55]. Also here our spectrally resolved algorithms can, thus, provide additional insights on the relation between brain rhythms and information-processing.

## Conclusion

In this study we have presented an algorithm that provides a spectral representation of the computational dynamics of neural processes in terms of the active information storage. Using this algorithm for the analysis of changes in neural information processing under anesthesia, we showed that this analysis can add valuable additional insights that are not provided by the analysis of changes in spectral power.

Our method is fully available and integrated in the open source package IDTXl:https://github.com/pwollstadt/IDTxl/tree/feature spectral ais, along with a demo script.

## Acknowledgements

We are grateful to K. Selller for the LFP ferrets recordings and F. Flöhlich to make the data available. MW is a professor at the Campus Institute for Dynamics of Biological Networks (CIDBN) funded by the Volkswagen Stiftung.

## References

1. Schreiber T. Measuring information transfer. Physical Review Letters. 2000;85(2):461–464. doi:10.1103/PhysRevLett.85.461.

2. Lizier JT. The local information dynamics of distributed computation in complex systems. Springer; 2013.

3. Lizier JT, Prokopenko M, Zomaya AY. Local measures of information storage in complex distributed computation. Information Sciences. 2012;208:39–54. doi:10.1016/j.ins.2012.04.016.

4. Lizier JT, Prokopenko M, Zomaya AY. Local information transfer as a spatiotemporal filter for complex systems. Physical Review E - Statistical, Nonlinear, and Soft Matter Physics. 2008;77(2):1–12. doi:10.1103/PhysRevE.77.026110.

5. Lizier JT, Flecker B, Williams PL. Towards a synergy-based approach to measuring information modification. IEEE Symposium on Artificial Life (ALIFE). 2013;2013-January(January):43–51. doi:10.1109/ALIFE.2013.6602430.

6. Gómez C, Lizier JT, Schaum M, Wollstadt P, Grützner C, Uhlhaas P, et al. Reduced predictable information in brain signals in autism pectrum disorder. Frontiers in Neuroinformatics. 2014;8(FEB):1–12. doi:10.3389/fninf.2014.00009.

7. Wibral M, Lizier JT, Vögler S, Priesemann V, Galuske R. Local active information storage as a tool to understand distributed neural information processing. Frontiers in Neuroinformatics. 2014;8(JAN):1–11. doi:10.3389/fninf.2014.00001.

8. Wollstadt P, Sellers KK, Rudelt L, Priesemann V, Hutt A, Fröhlich F, et al. Breakdown of local information processing may underlie isoflurane anesthesia effects. PLoS Computational Biology. 2017;13(6):1–35. doi:10.1371/journal.pcbi.1005511.

9. Wollstadt P, Rathbun DL, and WMU, Bastos AM, Lindner M, Priesemann V, et al. Information-theoretic analyses of neural data to minimize the effect of researchers’ assumptions in predictive coding studies. arXiv. 2022;.

10. Pinzuti E, Wollstadt P, Gutknecht A, Tüscher O, Wibral M. Measuring spectrally-resolved information transfer. PLoS Computational Biology. 2020;16(12 December):1–40. doi:10.1371/journal.pcbi.1008526.

11. Zipser D, Kehoe B, Littlewort G, Fuster J. A spiking network model of short-term active memory. Journal of Neuroscience. 1993;13(8):3406–3420. doi:10.1523/jneurosci.13-08-03406.1993.

12. Obst O, Boedecker J, Schmidt B, Asada M. On active information storage in input-driven systems. arXiv. 2013;(January 2015).

13. Brodski-Guerniero A, Paasch GF, Wollstadt P, Özdemir I, Lizier JT, Wibral M. Information-Theoretic Evidence for Predictive Coding in the Face-Processing System. The Journal of neuroscience : the official journal of the Society for Neuroscience. 2017;37(34):8273–8283. doi:10.1523/JNEUROSCI.0614-17.2017.

14. Brodski-Guerniero A, Naumer MJ, Moliadze V, Chan J, Althen H, Ferreira-Santos F, et al. Predictable information in neural signals during resting state is reduced in autism spectrum disorder. Human Brain Mapping. 2018;39(8):3227–3240. doi:10.1002/hbm.24072.

15. Sellers KK, Bennett DV, Hutt A, Fröhlich F. Anesthesia differentially modulates spontaneous network dynamics by cortical area and layer. Journal of Neurophysiology. 2013;110(12):2739–2751. doi:10.1152/jn.00404.2013.

16. Bastos AM, Lundqvist M, Waite AS, Kopell N, Miller EK. Layer and rhythm specificity for predictive routing. Proceedings of the National Academy of Sciences of the United States of America. 2020;117(49):31459–31469. doi:10.1073/pnas.2014868117.

17. Bastos AM, Vezoli J, Bosman CA, Schoffelen JM, Oostenveld R, Dowdall JR, et al. Visual areas exert feedforward and feedback influences through distinct frequency channels. Neuron. 2015;85(2):390–401. doi:10.1016/j.neuron.2014.12.018.

18. Purdon PL, Pierce ET, Mukamel EA, Prerau MJ, Walsh JL, Wong KFK, et al. Electroencephalogram signatures of loss and recovery of consciousness from propofol. Proceedings of the National Academy of Sciences of the United States of America. 2013;110(12). doi:10.1073/pnas.1221180110.

19. Percival DB, Walden AT. Wavelet methods for time-series analysis. Cambridge University Press; 2013.

20. Walden AT. Wavelet analysis of discrete time series. European Congress of Mathematics. 2001;202:627–641.

21. Wollstadt P, Lizier J, Vicente R, Finn C, Martinez-Zarzuela M, Mediano P, et al. IDTxl: The Information Dynamics Toolkit xl: a Python package for the efficient analysis of multivariate information dynamics in networks. Journal of Open Source Software. 2019;4(34):1081. doi:10.21105/joss.01081.

22. Wollstadt P, Hasenjäger M, Wiebel-Herboth CB. Quantifying the predictability of visual scanpaths using active information storage. Entropy. 2021;23(2):1–14. doi:10.3390/e23020167.

23. Wibral M, Lizier JT, Priesemann V. Bits from brains for biologically inspired computing. Frontiers in Robotics and AI. 2015;2:5.

24. Faes L, Nollo G, Porta A. Information-based detection of nonlinear Granger causality in multivariate processes via a nonuniform embedding technique. Physical Review E - Statistical, Nonlinear, and Soft Matter Physics. 2011;83:1–15. doi:10.1103/PhysRevE.83.051112.

25. Takens F. Detecting strange attractors in turbulence. Dynamical Systems and Turbulence, Lecture Notes in Mathematics. 1981;898:366–381. doi:10.1007/bfb0091924.

26. Lancaster G, Iatsenko D, Pidde A, Ticcinelli V, Stefanovska A. Surrogate data for hypothesis testing of physical systems. Physics Reports. 2018;748:1–60. doi:10.1016/j.physrep.2018.06.001.

27. Breakspear M, Brammer M, Robinson PA. Construction of multivariate surrogate sets from nonlinear data using the wavelet transform. Physica D: Nonlinear Phenomena. 2003;182(1-2):1–22. doi:10.1016/S0167-2789(03)00136-2.

28. Keylock CJ. Constrained surrogate time series with preservation of the mean and variance structure. Physical Review E - Statistical, Nonlinear, and Soft Matter Physics. 2006;73(3):2–5. doi:10.1103/PhysRevE.73.036707.

29. Keylock CJ. Characterizing the structure of nonlinear systems using gradual wavelet reconstruction. Nonlinear Processes in Geophysics. 2010;17(6):615–632. doi:10.5194/npg-17-615-2010.

30. Percival DB. Analysis of geophysical time series using discrete wavelet transforms: An overview. Lecture Notes in Earth Sciences. 2008;112:61–79. doi:10.1007/978-3-540-78938-34.

31. Cornish CR, Bretherton CS, Percival DB. Maximal overlap wavelet statistical analysis with application to atmospheric turbulence. Boundary-Layer Meteorology. 2006;119(2):339–374. doi:10.1007/s10546-005-9011-y.

32. Florin E, Gross J, Pfeifer J, Fink GR, Timmermann L. The effect of filtering on Granger causality based multivariate causality measures. NeuroImage. 2010;50(2):577–588. doi:10.1016/j.neuroimage.2009.12.050.

33. Zhang Z, Telesford QK, Giusti C, Lim KO, Bassett DS. Choosing wavelet methods, filters, and lengths for functional brain network construction. PLoS ONE. 2016;11(6). doi:10.1371/journal.pone.0157243.

34. Xiong. Entropy measures, entropy estimators, and their performance in quantifying complex dynamics: Effects of artifacts, nonstationarity, and long-range correlations. Physiology & behavior. 2017;176(5):139–148. doi:10.1103/PhysRevE.95.062114.Entropy.

35. Vakorin VA, Krakovska OA, McIntosh AR. Confounding effects of indirect connections on causality estimation. Journal of Neuroscience Methods. 2009;184(1):152–160. doi:10.1016/j.jneumeth.2009.07.014.

36. Salvatier J, Wiecki TV, Fonnesbeck C. Probabilistic programming in Python using PyMC3. PeerJ Computer Science. 2016;2:e55.

37. Vehtari A, Lampinen J. Bayesian model assessment and comparison using cross-validation predictive densities. Neural Computation. 2002;14(10):2439–2468. doi:10.1162/08997660260293292.

38. Gelman A, Carlin JB, Stern HS, Rubin DB. Bayesian Data Analysis; 2004.

39. Fries P. Communication Through Coherence (CTC 2.0). Neuron. 2015;88(1):220–235. doi:10.1016/j.neuron.2015.09.034.Rhythms.

40. Singer W. Neuronal oscillations: unavoidable and useful? European Journal of Neuroscience. 2018;48(7):2389–2398. doi:10.1111/ejn.13796.

41. Fröhlich F. Endogenous and exogenous electric fields as modifiers of brain activity: Rational design of noninvasive brain stimulation with transcranial alternating current stimulation. Dialogues in Clinical Neuroscience. 2014;16(1):93–102. doi:10.31887/dcns.2014.16.1/ffroehlich.

42. Franks NP. General anaesthesia: From molecular targets to neuronal pathways of sleep and arousal. Nature Reviews Neuroscience. 2008;9(5):370–386. doi:10.1038/nrn2372.

43. Bastos AM, Usrey WM, Adams RA, Mangun GR, Fries P, Friston KJ. Canonical Microcircuits for Predictive Coding. Neuron. 2012;76(4):695–711. doi:10.1016/j.neuron.2012.10.038.

44. Douglas RJ, Martin KAC. Neuronal circuits of the neocortex. Annual Review of Neuroscience. 2004;27(1):419–451. doi:10.1146/annurev.neuro.27.070203.144152.

45. Ferro D, van Kempen J, Boyd M, Panzeri S, Thiele A. Directed information exchange between cortical layers in macaque V1 and V4 and its modulation by selective attention. Proceedings of the National Academy of Sciences of the United States of America. 2021;118(12). doi:10.1073/pnas.2022097118.

46. Amigó JM, Monetti R, Tort-Colet N, Sanchez-Vives MV. Infragranular Layers Lead Information Flow during Slow Oscillations According to Information Directionality Indicators. J Comput Neurosci. 2015;39(1):53–62. doi:10.1007/s10827-015-0563-7.

47. Hudetz AG. Suppressing consciousness: Mechanisms of general anesthesia. Seminars in Anesthesia, Perioperative Medicine and Pain. 2006;25(4):196–204. doi:https://doi.org/10.1053/j.sane.2006.09.003.

48. Kihlstrom JF, Cork RC. Consciousness and Anesthesia. The Blackwell Companion to Consciousness. 2007;322(November):628–639. doi:10.1002/9780470751466.ch50.

49. Hagihira S. Changes in the electroencephalogram during anaesthesia and their physiological basis. British Journal of Anaesthesia. 2015;115:i27–i31. doi:10.1093/bja/aev212.

50. Weiner VS, Zhou DW, Kahali P, Stephen EP, Robert A. Propofol disrupts alpha dynamics in distinct thalamocortical networks underlying sensory and cognitive function during loss of consciousness. bioRxiv. 2022;.

51. Faes L, Pereira MA, Silva ME, Pernice R, Busacca A, Javorka M, et al. Multiscale information storage of linear long-range correlated stochastic processes. Physical Review E. 2019;99(3):1–13. doi:10.1103/PhysRevE.99.032115.

52. Costa M, Goldberger A, Peng CK. Multiscale Entropy Analysis of Complex Physiologic Time Series. Physical review letters. 2002;89:068102. doi:10.1103/PhysRevLett.89.068102.

53. Wibral M, Lizier JT, Vicente R. Directed Information Measures in Neuroscience. Springer; 2014.

54. Rudelt L, González Marx D, Wibral M, Priesemann V. Embedding optimization reveals long-lasting history dependence in neural spiking activity. PLOS Computational Biology. 2021;17(6):1–51. doi:10.1371/journal.pcbi.1008927.

55. Lundqvist M, Herman P, Warden MR, Brincat SL, Miller EK. Gamma and beta bursts during working memory readout suggest roles in its volitional control. Nature Communications. 2018;9(1):1–12. doi:10.1038/s41467-017-02791-8.

